# Agentic AI enables cell segmentation-free cell-type inference and cell-type-guided cell segmentation in spatial transcriptomics

**DOI:** 10.64898/2026.11.05.764509

**Authors:** Yuhui Jeong, Hanjae Lee, Donggeon Lee, Hyeon-Min Kim, So-Hui Kim, Suhyeon Park, Junseong Bae, Eunae Sandra Cho, Yoon-La Choi, Ohsang Kwon, Jong-Il Kim, Heetak Lee, Jeongbin Park

**Affiliations:** Department of Information Convergence Engineering, Pusan National University, Yangsan, Republic of Korea; Department of Dermatology, Seoul National University College of Medicine, Seoul, Korea; Genomic Medicine Institute (GMI), Medical Research Center, Seoul National University, Seoul, Korea; Laboratory of Cutaneous Aging and Hair Research, Clinical Research Institute, Seoul National University Hospital, Seoul, Korea; Lab of Molecular and Digital Pathology for Theranostics, Samsung Medical Center, Sungkyunkwan University School of Medicine, Seoul, Republic of Korea; Department of Digital Health, Samsung Advanced Institute for Health Sciences & Technology, Sungkyunkwan University, Seoul, Republic of Korea; Medical Imaging Insight Institute, Sungkyunkwan University School of Medicine, Seoul, Republic of Korea; Interdisciplinary Program of Genomic Data Science, Pusan National University, Yangsan, Republic of Korea; Department of Biological Sciences, Pusan National University, Pusan, Republic of Korea; Department of Oral Pathology, Oral Cancer Research Institute, BK21 FOUR Project, Yonsei University College of Dentistry, Seoul, Republic of Korea; Center for Genome Engineering, Institute for Basic Science, Daejeon, Republic of Korea; Department of Pathology and Translational Genomics, Samsung Medical Center, Sungkyunkwan University School of Medicine, Seoul, Republic of Korea; Department of Biochemistry and Molecular Biology, Seoul National University College of Medicine, Seoul, Korea; Center for Genome Engineering, Institute for Basic Science (IBS), Daejeon, Republic of Korea; Department of Biomedical Informatics, School of Medicine, Pusan National University, Yangsan, Republic of Korea; Department of Convergence Medical Science, School of Medicine, Pusan National University

## Abstract

Cell-type identification in spatial transcriptomics often relies on cell segmentation, which can introduce inaccuracies in cell boundaries and transcript assignment. Segmentation-free or cell-free methods have been developed to completely avoid cell segmentation, but inaccurate cell typing hinder accurate downstream analysis. Here, we introduce SSAF-T, a segmentation-free agentic framework that identifies cell types directly from spatial gene expression without predefined reference datasets. For the reference-free cell typing, SSAF-T incorporates CeLLMan, which employs expert-level agents for each organ to annotate cell types using a literature-grounded knowledge graph. We benchmark SSAF-T on both array-based and imaging-based spatial transcriptomics datasets in terms of accuracy, reproducibility, and scalability. We further show that SSAF-T provides an accurate cell segmentation based on the segmentation-free mapping, which ensure compatibility to existing downstream analysis ecosystem, and eventually enable quantitative analysis at single-cell level.

## Introduction

Spatial transcriptomics (ST) is revolutionizing the field as it can directly infer the cellular organization and interactions in a spatial context^1–3^. In terms of data analysis of ST datasets, cell segmentation is known as one big challenge for downstream analyses, such as clustering, cell-type annotation, niche inference, and interaction analysis^1,4,5^. Accurate cell segmentation remains a bottleneck, with reported limitations including missed nuclei, inaccurate cell boundaries, and unassigned transcripts^4–11^. Recently, segmentation-free methods, including SSAM^12^, FICTURE^13^, TopACT^14^, Sainsc^15^, and STHD^16^, have been developed to avoid cell segmentation for downstream analysis. These methods infer spatial cell-type distributions or molecular patterns from spatial gene expression without defining cell boundaries^12–16^.

However, such segmentation-free methods have two limitations. First, they require an external, matched scRNA-seq dataset or de novo clustering, both difficult to achieve^12–17^. For example, TopACT and STHD rely on external single-cell reference datasets, making them difficult to apply when suitable scRNA-seq reference data are limited^14,16,17^. SSAM, Sainsc, and FICTURE can work without an external reference, however, they require clustering or at least an estimation of the expected number of cell types in the dataset, which eventually requires cell typing, which is challenging^12,13,15,18^. Second, as segmentation-free analysis methods do not provide “cells”, they are not easily compatible with the widely used graph-based downstream analyses treating each cell as a node, such as Squidpy^19,20^. Especially, many studies focus on investigating direct cell-cell interactions between cells in proximity, but it is not intuitive to consider such analyses based on the outcome of such segmentation-free analysis^1,19,20^.

We suggest that these problems could be solved with an accurate, automated cell-type annotation algorithm that could be used in the spatial context, for both reference-free and segmentation-free ST data analysis. Not only can it infer the cell types accurately, assuming that such an algorithm can accurately identify the areas occupied by cell types, but cell segmentation could also be much improved using the areas as a prior, rather than solely relying on images such as DAPI, Poly-A, H&E, etc., which do not provide accurate boundaries between different cell types^4–8,10,14^.

In this regard, we considered using recently developed LLM-based automatic cell-typing methods to automate the cell-type annotation process in combination with our previous segmentation-free framework, SSAM^12,21–24^. However, unfortunately, it is known that the performance of LLM-based cell-typing still does not reach the level of human experts, and expert-driven manual curation based on marker-gene expression remains important for accurate cell-type annotation^21–25^. Moreover, SSAM was developed before the rise of agentic workflows in biosciences, and therefore there was a strong need for a new framework that natively supports LLMs. Thus, we ended up developing a novel framework that integrates LLM-based, automatic cell-type annotation with marker-gene evidence to achieve segmentation-free analysis that supports any spatial dataset regardless of its underlying technology.

To this end, here we introduce SSAF-T (<u>S</u>patial <u>S</u>egmentation-free <u>A</u>gentic <u>F</u>ramework for <u>T</u>ranscriptomics). SSAF-T identifies cell types without prior cell segmentation or predefined reference datasets, while allowing human experts to guide and refine the annotation process when needed. The key component of SSAF-T is its annotation agent, CeLLMan, which uses a curated, literature-grounded knowledge graph to assign standardized cell-type labels based on marker-gene evidence, improving annotation accuracy compared with an LLM without access to this knowledge. SSAF-T also performs a 2D or 3D cell segmentation by default, using the segmentation-free cell typing result as a prior, to ensure compatibility with any downstream analysis pipelines.

In this paper, we demonstrate the performance and applicability of SSAF-T and CeLLMan using publicly available and novel datasets as follows: (1) validating the cell-typing accuracy of CeLLMan against reference annotations using two scRNA-seq atlases (Tabula Sapiens and Tabula Muris Senis), (2) validating the performance of SSAF-T on six spatial transcriptomics datasets (Visium HD mouse small intestine, Stereo-seq mouse brain, Xenium mouse kidney, MERFISH mouse liver, Xenium human placenta, and Xenium human heart), and (3) applying SSAF-T to a newly generated Xenium mouse skin dataset for a discovery of previously unreported biological finding. We show that CeLLMan identifies cell types beyond published atlas annotations and that SSAF-T resolves spatial cell-type distributions consistent with tissue anatomy, while recovering transcripts missed by conventional segmentation methods.

## Results

### The SSAF-T computational framework

SSAF-T use SpatialData^26^ object as input and output, enabling analysis within existing spatial transcriptomics workflows that supports Scverse^27^ ecosystem. The framework consists of 6 steps (Fig. 1A).

**Figure 1.**
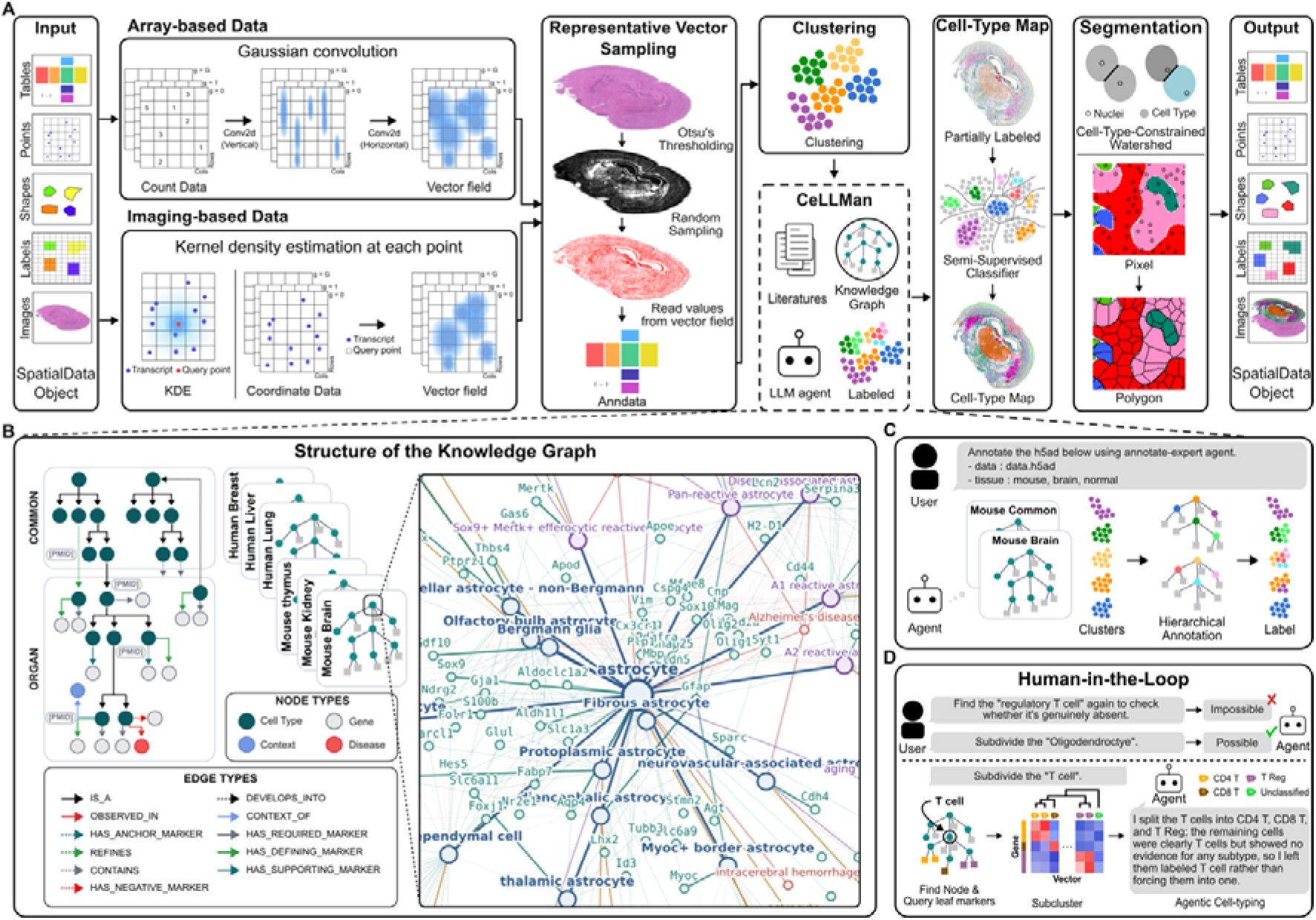
SSAF-T overview. (A) SSAF-T workflow. SSAF-T uses a SpatialData object as input and output. Step 1 converts spatial gene expression into a gene-expression vector field: array-based gene counts on a spatial grid through Gaussian convolution, and imaging-based transcript coordinates through kernel density estimation (KDE). Step 2 samples representative expression vectors within a tissue mask generated using Otsu thresholding of the staining image. Step 3 clusters the sampled vectors. Step 4 annotates each cluster using CeLLMan, an LLM-based annotation agent supported by a literature-grounded cell-type knowledge graph. Step 5 uses the annotated expression vectors to train a classifier that predicts cell-type labels across the entire vector field, generating a spatial cell-type map. Step 6 performs cell segmentation using a nucleus-seeded watershed constrained by the cell-type map. (B) Structure of the knowledge graph, organized into species-organ-specific modules, with cell types linked to marker genes supported by PubMed references (PMIDs). (C) An example “Default Annotation”. The agent selects the corresponding knowledge-graph module, evaluates each cluster’s profile, and assigns cell-type labels hierarchically. (D) Human-in-the-loop annotation. User requests to refine cell-type annotations are accepted or rejected based on the marker-gene evidence.

First, spatial gene expression data are either converted into continuous gene expression density matrices (imaging-based data), or Gaussian kernel convoluted (array-based data). For imaging-based data, transcript coordinates are used to estimate the spatial expression density of each gene through kernel density estimation (KDE) with a Gaussian kernel. For array-based data, we apply Gaussian convolution to account for noises, such as the one resulted by lateral diffusion of molecules. As previously explored in SSAM, the resulting gene-specific expression distributions are stacked to construct a multichannel “gene expression vector field”, in which a vector of estimated gene expression values represents each spatial position at a pixel or voxel.

Second, the gene expression vector field is downsampled by selecting representative positions within tissue regions likely to contain cells. To identify these regions, tissue staining images, such as H&E or morphology images, are processed using Otsu thresholding^28^ to generate a binary tissue mask. Spatial positions are then randomly sampled within the mask, and the corresponding gene expression vectors are extracted from the field to construct an expression matrix.

Third, the sampled expression vectors are clustered based on their gene expression profiles. Here we used Leiden^29^ clustering algorithm, followed by L1 normalization, log transform, and principal component analysis (PCA). To accelerate this process, we harnessed Rapids Single Cell^30^ library for GPU-accelerated computation for a scalable data analysis.

Fourth, each cluster is annotated by CeLLMan, an agentic cell-type annotation framework that uses a curated, literature-grounded knowledge graph (KG) of cell types and their marker genes (Fig. 1C, Supplementary Fig. 1). Here, the per-species knowledge graph contains organ modules, which contains cell type labels, marker genes, and spatial/disease context as nodes, and literatures that supports the nodes as evidence selected from PubMed database across the entire publication records in history as edges (Supplementary Fig. 3B). The cell types shared across multiple organs stored in a common module to avoid redundancy (Fig. 1B).

Within the KG, CeLLMan selects the appropriate module based on tissue information (Supplementary Fig. 1A), evaluates the marker-gene expression profile of each cluster against the knowledge graph (Supplementary Fig. 1B), and assigns a cell-type label. Annotation proceeds through iterative rounds of hierarchical classification, from broad cell classes to more specific cell types. In each round, three independent agents evaluate the marker-gene evidence and propose cell-type assignments, with the consensus decision determining whether and how annotation proceeds to be narrower or broader. When CeLLMan consensus decision reaches a conclusion that a cluster potentially contains multiple cell types, CeLLMan performs subclustering using subtype-specific marker genes and re-evaluates the resulting subclusters. Subclusters with different identities are annotated separately, whereas those sharing the same identity are merged into a single cluster.

By default, CeLLMan performs “Default Annotation”, which automatically repeats the above process iteratively until no further refinement is supported by discriminative marker-gene expression. In the end, the clusters lacking sufficient evidence for a specific cell type retain their parent labels rather than being assigned as cell types with no supporting evidence (Fig. 1C, Supplementary Fig. 1C). After this procedure, CeLLMan provides a human-in-the-loop step that allows further refining of cell-type annotations via natural-language instructions (Fig. 1D). Upon user requests in natural language, the agent either accepts further fine-subclustering of a cell type supported by evdences in KG, or rejecting the request if insufficient evidence is available.

Fifth, the sampled gene expression vectors and their cell-type labels assigned using CeLLMan are used to construct cell-type-specific expression profiles. We incorporate Latent Dirichlet Allocation (LDA), a probabilistic model to expand cell type labels to the entire vector field, using the downsampled vectors as the support, which eventually generates a fully annotated spatial cell-type map at every pixel/voxel location (details in Methods). As also explored in SSAM, the segmentation-free cell-type map can be used for segmentation-free downstream analyses, including spatial adjacency and tissue composition analyses (Supplementary Figs. S5-6).

Sixth, SSAF-T performs cell segmentation using the generated cell-type map and nuclei markers. Here we used watershed algorithm^31^ restricted by the cell-type boundaries defined in the segmentation-free cell-type map, using nuclei markers as seeds to identify individual cells. In the case of the cells having the same cell type in proximity are separated at the midpoint between neighboring nuclei. Here, cell segmentation can be partially performed to exclude certain cell types, such as multinucleated cells which is known to be extremely difficult to segment, preserving complex cellular structures that are difficult to segment while segmenting other cell types into individual cells (Supplementary Figs. S5-6).

Across the whole analysis steps, all outputs are stored in a SpatialData object, with the cell-type map stored in images and labels, segmentation polygons in shapes, and cell-level transcript counts and CeLLMan annotation results for the downsampled expression vectors in tables.

### CeLLMan complements published cell-type annotations in single-cell atlases

To evaluate the cell-type annotation performance of CeLLMan, we benchmarked CeLLMan with Tabula Sapiens v2.0^32^ (29 organs) and Tabula Muris Senis^33^ (18 organs) datasets. We compared the Default Annotation by CeLLMan against the published atlas annotations, using the marker gene expressions as the evidence of their cell-type identities. Here, each cell was classified into one of the following five categories: the same identity (identical), CeLLMan identified more detailed sub-cell types (CeLLMan narrower), reference reported more detailed sub-cell types (CeLLMan broader), conflict in cellular lineages (conflict), or either CeLLMan or reference reported as an unknown cell type (declined) (Supplementary Fig. 2A). As a result, 37.9% of cells were classified as identical, 37.2% as CeLLMan narrower, 9.4% as CeLLMan broader, and 15.3% as conflict for Tabula Sapiens dataset. Also, the corresponding percentages were 28.7%, 45.9%, 2.6%, and 22.6%, respectively for Tabula Muris Senis dataset (Supplementary Fig. 2A). The number of distinct cell-type labels was 797 for CeLLMan and 702 for the published annotations in Tabula Sapiens, and 447 and 221, respectively, in Tabula Muris Senis (Supplementary Fig. 2B).

First, we evaluated whether CeLLMan’s narrower/broader cell-type annotations were supported by marker-gene expression. Here, we assessed whether markers specific to the narrower labels distinguished the corresponding cells from other cells assigned the same broader cell type. Among cells classified as CeLLMan narrower, marker-gene evidence supported CeLLMan in 88.0% of Tabula Sapiens cells and 74.5% of Tabula Muris Senis cells, whereas only 1.0% and 13.3% supported the published annotations, respectively (Supplementary Fig. 2C). Among cells classified as CeLLMan broader, marker-gene evidence supported CeLLMan in 39.0% of Tabula Sapiens and 44.3% of Tabula Muris Senis cells, whereas only 13.5% and 17.9% supported the published annotations, respectively (Supplementary Fig. 2D).

Next, we determined which annotation better supports the marker-gene expression for cells classified as conflict. We compared the average gene expression of the marker genes (marker-gene scores) using cells assigned the same cell type by both CeLLMan and the published annotations as references for each competing cell type. Marker-gene evidence supported CeLLMan in 56.2% of Tabula Sapiens cells and 61.1% of Tabula Muris Senis cells, whereas the published annotations supported it in 7.4% and 2.3%, respectively (Supplementary Fig. 2E).

Finally, we combined the marker-based evaluation results across the CeLLMan narrower, CeLLMan broader, and conflict categories for each organ and calculated the proportions of cells supporting either annotation (Supplementary Fig. 2F). Across all organs, marker-gene evidence supported CeLLMan in 45.0% of Tabula Sapiens cells and 49.1% of Tabula Muris Senis cells, compared with 2.8% and 7.1%, respectively, supporting the published annotations.

These results show that CeLLMan overall better identifies the actual cell-type identities compared to the previously published cell-type annotations for Tabula Sapiens / Tabula Muris Senis datasets, which indicates a potential to complement manual annotations by experts in identifying cell-types within such large datasets.

### SSAF-T reconstructs tissue architecture in array-based spatial transcriptomics data

To evaluate the performance of SSAF-T for analyzing array-based spatial transcriptomics data, we applied SSAF-T to two representative datasets: a Visium HD mouse small intestine dataset from Kamel *et al*.^34^ and a Stereo-seq mouse brain dataset from Chen *et al.*^35^. The resulting cell-type maps and segmented cells were compared with the published results (Fig. 2A, B). In the small intestine, ENACT algorithm was used to annotate cell types, where most cells in the enlarged villus region as B cells. In contrast, SSAF-T distinguished enterocytes, plasma cells, macrophages, goblet cells, and Paneth cells which better supports the known biology. In the brain, SSAF-T distinguished anatomically distinct cell types in the neocortex, CA1, and amygdala that were broadly annotated as glutamatergic or hippocampal pyramidal neurons in the published analysis. These cell-type assignments were consistent with the spatial distributions of their marker genes (Fig. 2B).

**Figure 2.**
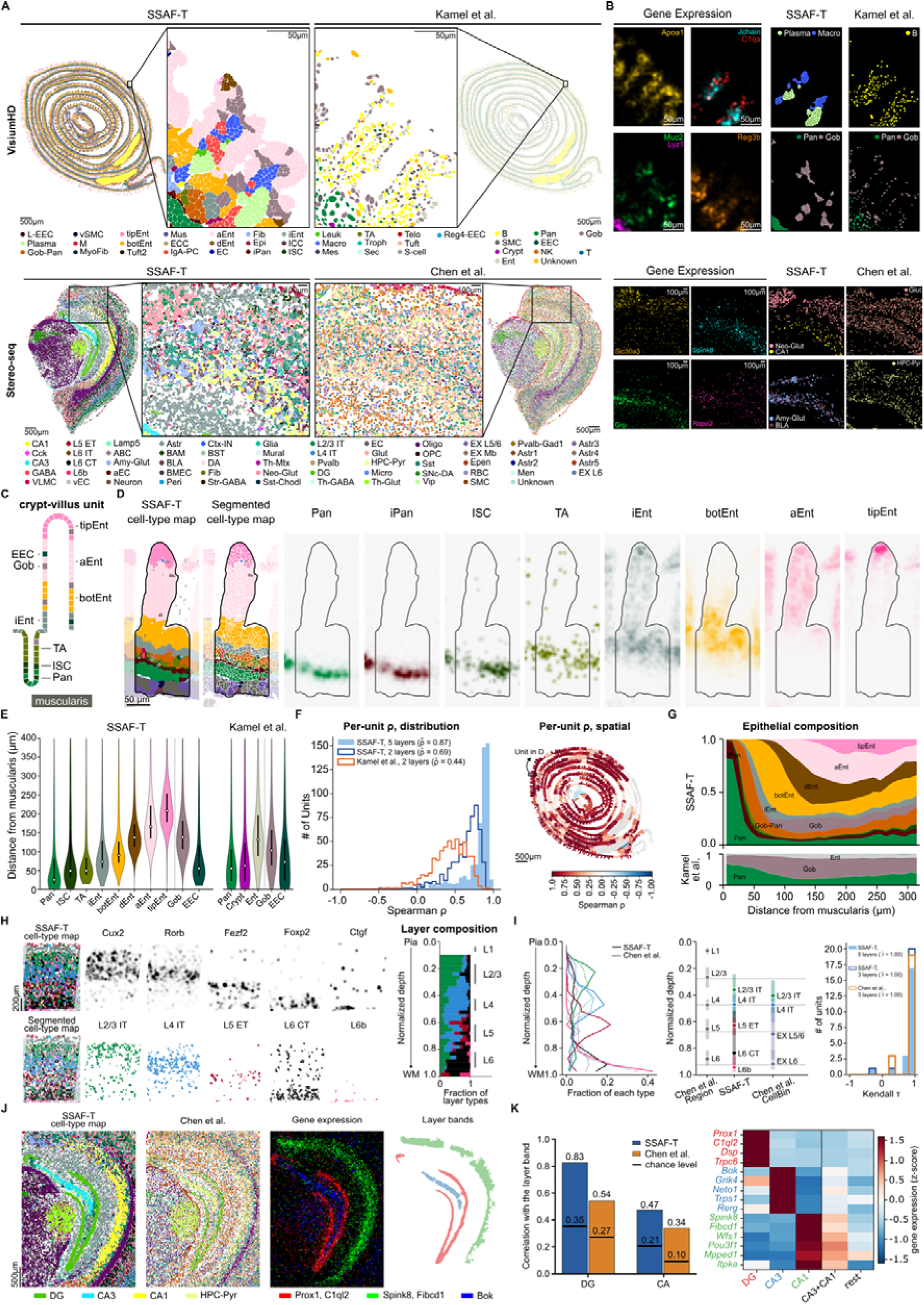
SSAF-T reconstructs tissue architecture in array-based spatial transcriptomics data. (A) Segmented cell-type maps of Visium HD mouse small intestine (top) and Stereo-seq mouse brain (bottom) generated by SSAF-T and the published analyses of the same sections: Kamel *et al.* using ENACT segmentation and Chen *et al.* using CellBin segmentation. Boxed regions are enlarged in the middle. (B) Gene-expression vector fields of the indicated marker genes and corresponding cell-type annotations in the enlarged regions. (C) Schematic of a crypt-villus unit showing the expected spatial order of epithelial cell types. (D) Representative crypt-villus unit showing the SSAF-T cell-type map, segmented cells, and gene-expression vector fields for eight epithelial cell types: Paneth cells (*Lyz1*, *Mmp7*), immature Paneth cells (*Olfm4*, *Lgr5*, *Lyz1*), intestinal stem cells (*Lgr5*, *Olfm4*, *Ascl2*, *Smoc2*), transit-amplifying cells (*Sox9*, *Mki67*, *Pcna*), immature enterocytes (*Fgfbp1*, *Prss32*, *Krt8*, *Slc16a1*), villus-bottom enterocytes (*Reg3b*, *Reg3g*, *Saa1*, *Ccl25*), absorptive enterocytes (*Alpi*, *Apoa1*, *Apoa4*, *Fabp1*, *Apob*, *Rbp2*, *Slc5a1*, *Anpep*, *Aldob*, *Krt20*, *Maob*, *Fabp2*, *Cdhr2*), and villus-tip enterocytes (*Nt5e*, *Ada*, *Slc28a2*, *Apoa4*, *Cdhr2*). (E) Distributions of epithelial cell distances from the muscularis in the representative crypt-villus unit, grouped by cell type. White points indicate medians, and bars indicate interquartile ranges. (F) Spearman’s correlation coefficients between epithelial cell distance from the muscularis and expected cell-type rank across crypt-villus units. Left, distributions at five-level (Pan/ISC, TA, iEnt, botEnt, and tipEnt) and matched two-level annotation resolutions (crypt: Pan/ISC/TA; absorptive: all enterocyte subtypes for SSAF-T; crypt: Pan/Crypt; absorptive: Ent for ENACT). Right, correlations for individual units, the box marking the unit shown in (D). (G) Epithelial cell-type composition across distances from the muscularis in SSAF-T and the published annotation. (H) Cortical organization in three adjacent units, showing the SSAF-T cell-type map, segmented cells, layer-specific marker-gene expression vector fields, and cell-type composition along normalized cortical depth. (I) Cortical layer distributions and spatial organization across normalized depth. Left, fractions of cortical layer cell types in SSAF-T (solid lines) and Chen *et al.* (dotted lines). Middle, median and interquartile range of normalized depth for each layer type in SSAF-T and the published CellBin-based and bin50 anatomical annotations. Right, distributions of Kendall’s τ between observed and expected layer order across cortical units at five-level (L2/3 IT, L4 IT, L5 ET, L6 IT/L6 CT, and L6b) and matched three-level resolutions (L2/3 IT, L4 IT, and L5 ET/L6 CT/L6b for SSAF-T; EX L2/3, EX L4, and EX L5/6/EX L6 for Chen *et al.*). (J) Enlarged cell-type maps of the hippocampal region from SSAF-T and Chen *et al.*, gene-expression vector fields of the indicated subtype markers, and layer bands defined by marker-gene expression. (K) Left, Pearson’s correlations between each annotation and the corresponding layer bands (DG and CA, with CA1 and CA3 combined for SSAF-T); horizontal lines indicate chance levels. Right, marker-gene expression across annotated cell types, z-scored across columns.

To assess whether SSAF-T recovered intestinal epithelial zonation, we examined cell-type distributions in detail along the crypt-villus axis (Fig. 2C-G). In a representative crypt-villus unit, the SSAF-T cell-type map was consistent with the expected anatomical organization and the spatial distributions of corresponding marker genes (Fig. 2C, D). Within this unit, the distributions of epithelial cell types by distance from the muscularis showed that SSAF-T distinguished distinct epithelial cell types, from Paneth cells and intestinal stem cells at the crypt base to progressively differentiated enterocytes toward the villus tip, whereas ENACT identified broader Paneth, crypt, and enterocyte categories (Fig. 2E). To quantify this organization across the tissue, we calculated Spearman’s correlation between epithelial cell position and the expected cell-type rank for each crypt-villus unit. At the five-level annotation resolution, the median correlation coefficient was 0.87 across 529 units. When both annotations were matched at a common two-level resolution, SSAF-T achieved a higher median correlation than ENACT (0.69 versus 0.44; Fig. 2F). Epithelial cell-type composition across the tissue also showed a gradual transition from crypt-associated cell types to differentiated enterocytes along the crypt-villus axis (Fig. 2G). These results indicate that SSAF-T resolves finer intestinal epithelial zonation while maintaining more consistent spatial organization at matched annotation resolution.

In the Stereo-seq mouse brain dataset, we examined cortical lamination and hippocampal organization (Fig. 2H-K). In representative cortical units, the SSAF-T cell-type map distinguished five neocortical layer cell types arranged in the expected order from the pia to the white matter, consistent with the spatial distributions of corresponding marker genes (Fig. 2H). This spatial organization was also consistent with the published bin50 anatomical regions, as shown by the cortical depth distributions of both SSAF-T and the published result. Kendall’s correlation between the observed and expected layer order reached a median of 1.00 for both methods across 22 cortical units (Fig. 2I). In the hippocampus, SSAF-T distinguished dentate gyrus, CA1, and CA3 cell types, with spatial distributions consistent with the layer bands (Fig. 2J). To quantify this correspondence, we calculated correlations between the cell-type annotations and the layer bands, combining CA1 and CA3 into a single CA region for comparison with the published annotation. SSAF-T showed higher correlations than the published result for both the dentate gyrus (0.83 versus 0.54) and CA (0.47 versus 0.34; Fig. 2K). Marker-gene expression further supported the distinction between CA1 and CA3 cell types identified by SSAF-T (Fig. 2K).

Together, these results demonstrate that SSAF-T reconstructs established tissue architecture in array-based spatial transcriptomics datasets, providing finer cell types while preserving the spatial organization of intestinal epithelial and neuronal cell types.

### SSAF-T improves cell segmentation in imaging-based spatial transcriptomics

To evaluate the performance of SSAF-T for analysis of imaging-based spatial transcriptomics data, we applied it to a Xenium mouse kidney dataset from Xuanyuan *et al.*^36^ and a MERFISH mouse liver dataset from Liu *et al.*^37^. The resulting cell-type maps and segmented cells were compared with the published results (Fig. 3A, B). In the kidney, the published annotation identified TAL and DCT as single cell types, whereas SSAF-T further distinguished TAL and cortical TAL, and DCT and DCT2. In the liver, SSAF-T distinguished periportal hepatocytes and pericentral hepatocytes along the zonation axis defined by *Pck1* and *Gulo* expression, respectively, while retaining hepatocytes in regions where these expression patterns overlapped. These cell-type distributions were consistent with the corresponding spatial marker-gene expressions (Fig. 3B).

**Figure 3.**
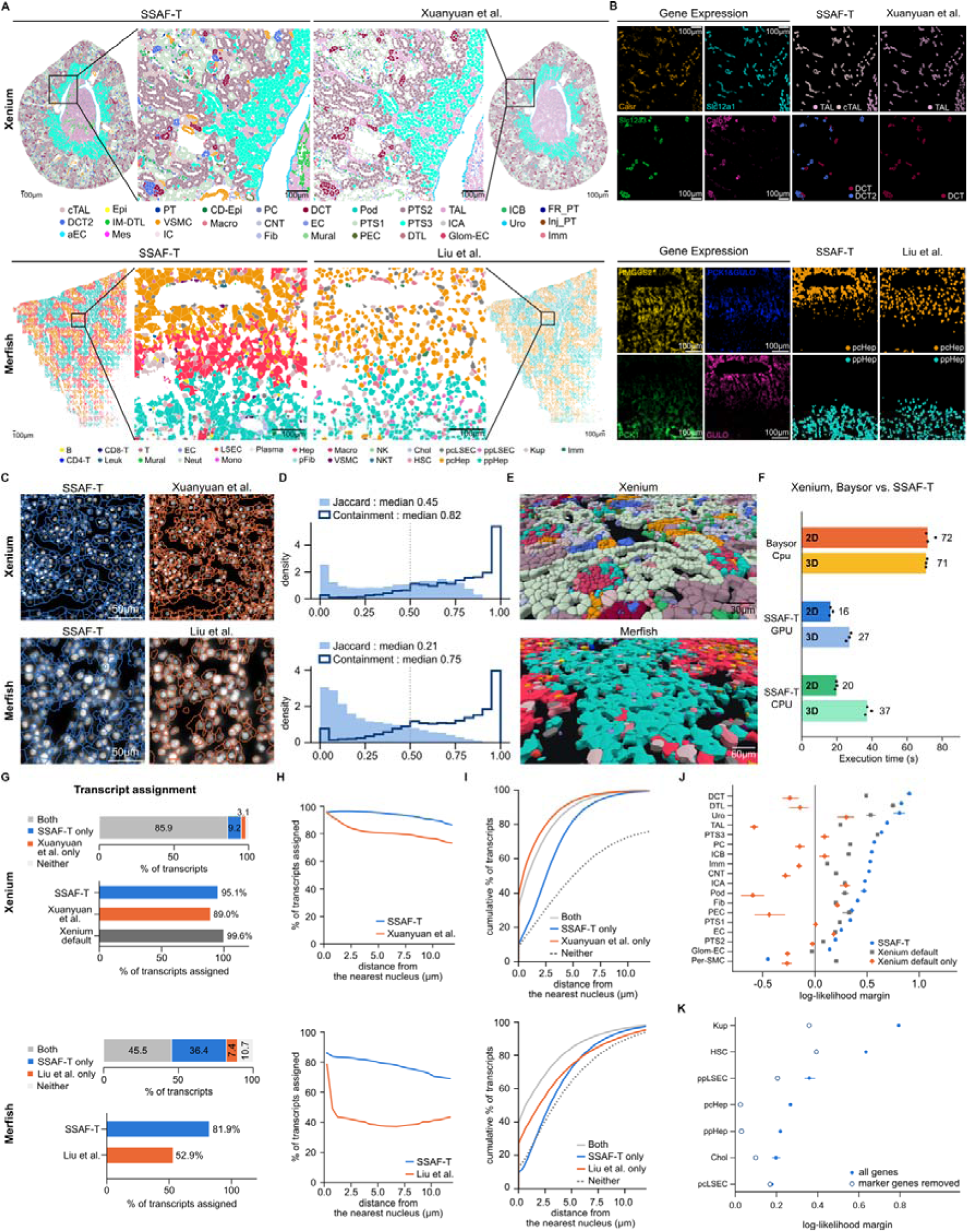
SSAF-T improves cell segmentation in imaging-based spatial transcriptomics data. (A) Segmented cell-type maps of Xenium mouse kidney (top) and MERFISH mouse liver (bottom) generated by SSAF-T and the published analyses of the same sections: Xuanyuan *et al.* using Baysor segmentation and Liu *et al.* using Cellpose segmentation. Boxed regions are enlarged in the middle. (B) Gene-expression vector fields of the indicated marker genes and corresponding cell-type annotations in the enlarged regions. (C) Cell boundaries from SSAF-T (blue) and the published segmentations (orange) overlaid on DAPI images. (D) Distributions of transcript containment and Jaccard indices between published cells and their best-matching SSAF-T cells. (E) Three-dimensional segmented cell-type maps generated by SSAF-T. (F) Execution times for 2D (top) and 3D (bottom) segmentation on the same Xenium kidney region. Bars indicate medians of three runs, and points indicate individual runs. (G) Transcript assignment rates in SSAF-T and the published segmentations. Top, proportions of transcripts assigned by both methods, SSAF-T only, the published segmentation only, or neither. (H) Transcript assignment rates as a function of distance to the nearest nucleus. (I) Empirical cumulative distribution functions (ECDFs) of distance to the nearest nucleus for the four transcript groups defined in (G). (J) Multinomial log-likelihood margins for transcripts excluded by Baysor but recovered by SSAF-T, Xenium default segmentation, or Xenium default segmentation alone. Error bars indicate 95% confidence intervals from 200 multinomial bootstrap replicates. (K) Multinomial log-likelihood margins for transcripts excluded by the published Cellpose segmentation but recovered by SSAF-T in MERFISH liver, calculated using all genes and after excluding knowledge-graph marker genes associated with the assigned cell type. Error bars indicate 95% confidence intervals from 300 multinomial bootstrap replicates.

Next, we tested the cell segmentation of SSAF-T against other cell segmentation algorithms. Here, we compared SSAF-T cell boundaries and transcript assignments with those obtained using transcript-based Baysor^38^ in Xenium (with Xenium cell segmentation as prior) and image-based Cellpose^39^ in MERFISH (Fig. 3C, D). We found that both published and SSAF-T cell segmentations agree in sharing most transcripts. The median transcript containment was 0.82 for Xenium and 0.75 for MERFISH, with corresponding median Jaccard indices of 0.45 and 0.21, respectively (Fig. 3D). We further examined the disagreements between the two methods by classifying transcripts into four categories, according to which cell they were originally assigned: SSAF-T, the published segmentation, both, or neither (Fig. 3G). SSAF-T assigned 95.1% of transcripts in Xenium and 81.9% in MERFISH, compared with 89.0% and 52.9% for the published segmentations, respectively. Transcript assignment rates decreased by the distance from the nearest nucleus – compared to Baysor and Cellpose, SSAF-T maintained higher assignment rates farther from the nucleus (Fig. 3H). Consistent with this pattern, transcripts recovered only by SSAF-T tend to exist farther from nuclei than those assigned by both methods, although the amount of extent is minute (Fig. 3I).

Therefore, we further evaluated whether these transcripts, abandoned by both methods but included by SSAF-T, are actual signals from a cell or background noise. For this, we evaluated their multinomial log-likelihood margins against reference expression profiles derived from the published annotations (Fig. 3J, K). In the Xenium kidney dataset, transcripts excluded by Baysor but recovered by SSAF-T showed positive margins for most evaluated cell types, generally exceeding those recovered by Xenium default segmentation. In contrast, transcripts assigned by Xenium default segmentation but excluded by both Baysor and SSAF-T showed negative margins for several cell types (Fig. 3J). These results suggest that SSAF-T recovers cell-type-associated transcripts beyond the Baysor segmentation boundaries, whereas transcripts additionally assigned by Xenium default segmentation show weaker cell-type-specific expression signals. In the MERFISH liver dataset, transcripts excluded by the published Cellpose segmentation but recovered by SSAF-T showed positive log-likelihood margins across all seven evaluated cell types (Fig. 3K).

Finally, in terms of running speed, SSAF-T completed both 2D and 3D segmentation faster than Baysor, even when running on a CPU (Fig. 3F).

### SSAF-T maps follicular cell identities and Satb2 expression

In a mouse skin wound healing model (Fig. 4A, Supplementary Fig. 7), we were able to use SSAF-T to identify hair follicle cell types with novel *Satb2* gene expression patterns in early to mid-anagen mouse hair follicles. The cell type, primarily labelled as ‘mesenchymal cell’, was located as a collar surrounding the matrix progenitors in only hair follicles adjacent to the wound area of post-wounding day 15 (Fig. 4B-C). This ‘mesenchymal cell’ labelled cell type was not seen in other hair follicles or tissue areas within the section, having a low confidence annotated by ceLLMan (Fig. 4D), and was Pdgrfa– and Acta2-, excluding the possibility of being true mesenchymal stromal cell types during a human-in-the-loop inspection by a human expert. Rather the cell type was suspected to be a set of two distinct cell types, a Maf+, Msx2+ inner root sheath (IRS) cell layer with an unexpected high Satb2+ expression, and a Lef1+, Msx2+ precortex layer, which spatial distribution well matched the known anagen stage hair follicle architecture (Fig 4E-G). Despite being primarily grouped together, Wnt5a expression was noticed mainly in the Satb2+ IRS layer and was minimal in the Satb2-precortex layer, implying finer cell subtype classification within the SSAF-T recognized ‘mesenchymal’ cell type layer (Fig 4H). The anagen hair follicle was composed of a Wnt5a+/Lef1+ and weak Satb2 matrix cells, followed by a Lef1+/Wnt5a-/Satb2-precortex and shaft cells, then Wnt5a+/Lef1– and Satb2 high IRS cells, with the latter two cell types being classified as a distinct cell type layer by SSAF-T annotation.

**Figure 4.**
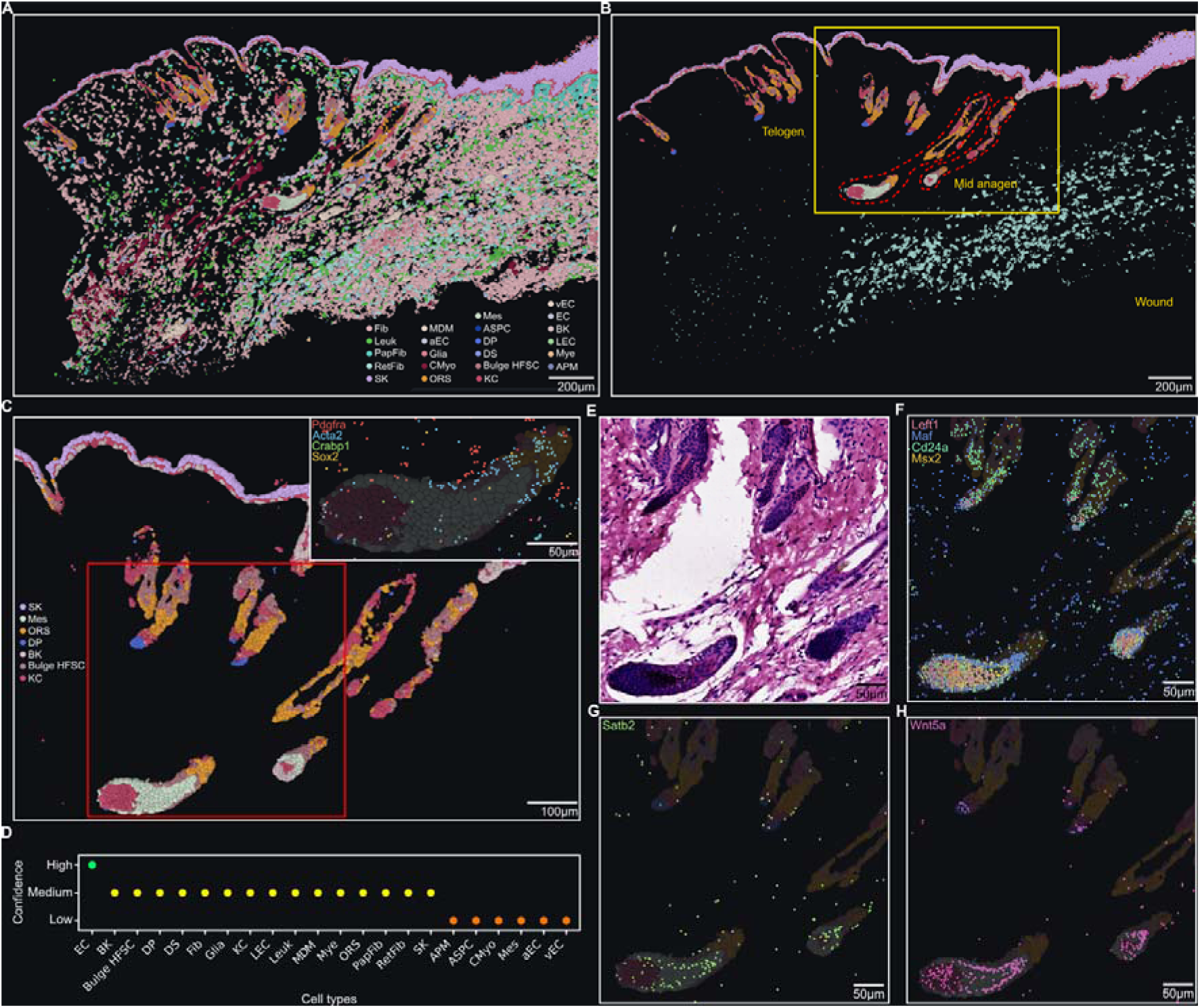
Mouse skin wound model (post 15 days) annotated with the SSAF-T framework reveals potential for novel gene expression or cell type recognition. (A) Overall spatial transcriptomic data and cell type annotation image of the mouse skin wound sample by the SSAF-T framework. Scale bars are inserted within its regarding image panel. (B) Injury-induced cycling and telogen-to-anagen transition seen in hair follicles adjacent to the wound. Dense reticular fibroblast populations are seen throughout the granulation tissue of the healing wound, while two anagen stage (red dotted line) and multiple telogen stage hair follicles are seen left to the healing wound. (C) Magnified view of the yellow-boxed region in (B). A light green colored cell layer labelled as ‘mesenchymal cell’ is uniquely seen between the epithelial components of the mid-anagen phase hair follicles next to the wound. (Inset) Pdgfra-, Acta2-, Sox2– and Crabp1-confirms the unique cell layer to be of epithelial origin, rather than being true mesenchymal cell types or dermal papilla. (D) Dot plot showing the confidence scores of CeLLMan annotations for each cell type. (E) H&E-stained image of the region indicated by the red box in (C). (F) The cell type architectures revealed by spatial transcriptomic patterns in anagen follicles. (Cd24a+/Lef1+ matrix cells, Lef1+/Msx2+ precortex cells, Maf+/Msx2+ inner root sheath cells). (G) Satb2 expression is seen in the peripheral of the unique cell layer, while minimally expressed in matrix cells and telogen hair follicle cell types, indicating possible heterogeneity in the unique cell layer populations. (H) Wnt5a (non-canonical Wnt pathway) along with Satb2 and Lef1 (canonical Wnt pathway) spatial expression patterns suggest distinct transcriptomic profiles of canonical and non-canonical Wnt pathway balance among anagen hair follicle cell types. The SSAF-T framework supported cell subtype recognition and annotation at a coarse resolution that correlated with specific gene expression patterns that have not yet been reported in previous literatures.

Interestingly, the three cell type gene profiles are all known to either directly or indirectly mediate the Wnt pathway. Both anagen and telogen hair follicles express noncanonical Wnt5a at cell types at the end of the hair follicle (e.g. bulge, hair germ, matrix cells and dermal papilla, depending on cycle phase), but the required balance between canonical and non-canonical Wnt pathway is cell type specific even within a single hair follicle^40,41^. Wnt5a has been reported to inhibit telogen-to-anagen transition during normal hair follicle cycling contributing to its appropriate structural polarity which well matched the IRS expressing Wnt5a pattern as seen in our data^42^. Satb2, a nuclear matrix attachment region binding transcription factor^43^, is known to mediate the canonical Wnt/β-catenin pathway in normal colon and gastrointestinal cancer^44–46^. Although the role of Satb2 in hair follicle Wnt pathway regulation has not been reported, elevated Satb2 expression in pilomatrix tumors has been reported, clinically supporting our observed Satb2 expression pattern to be a valid biological finding^47^. As the spatial transcription of core Wnt pathway mediators, Satb2, Wnt5a, Lef1 and Wls (data not shown), are distinct among hair follicle cell types in our mouse skin sample, we carefully suggest that the SSAF-T cell annotation framework showed potential in discriminating unique gene expression patterns and cell type candidates as a separate annotation layer in Xenium generated datasets. The detailed relation of Satb2, Wnt pathway mediators and hair follicle cell types is out of the scope of this paper, so relevant quantification or experimental validation was not done on the subject.

### SSAF-T identifies and maps macrophages in lung cancer across different spatial locations

Macrophages arise from two lineages. Tissue-resident macrophages (TRMs), which reside in the alveoli, self-renew locally. Monocyte-derived macrophages (MDMs) originate from adult hematopoietic stem cells and accumulate in inflammatory lesions^48^. In non-small cell lung cancer (NSCLC), TRMs are depleted in tumors relative to normal lung and are redistributed to the tumor periphery as the tumor grows, so that the tumor microenvironment becomes dominated by MDMs^48^.

An annotation method that correctly distinguishes these two populations should therefore recapitulate this shift in composition. Specifically, the proportion of TRMs among the two populations should be highest in normal lung, intermediate at the tumor border, and lowest in the tumor core. Using this gradient as a benchmark, we evaluated whether SSAF-T distinguish TRMs from MDMs better than a conventional approach. We defined three regions on a Xenium Prime section of human lung adenocarcinoma: normal lung, tumor border, and tumor core (Fig. 5A). In each region, we compared SSAF-T cells annotated with CeLLMan against cells from our in-house pipeline. The in-house pipeline annotates the default Xenium segmentation with SingleR, using a lung cancer atlas as the reference^49,50^.

**Figure 5.**
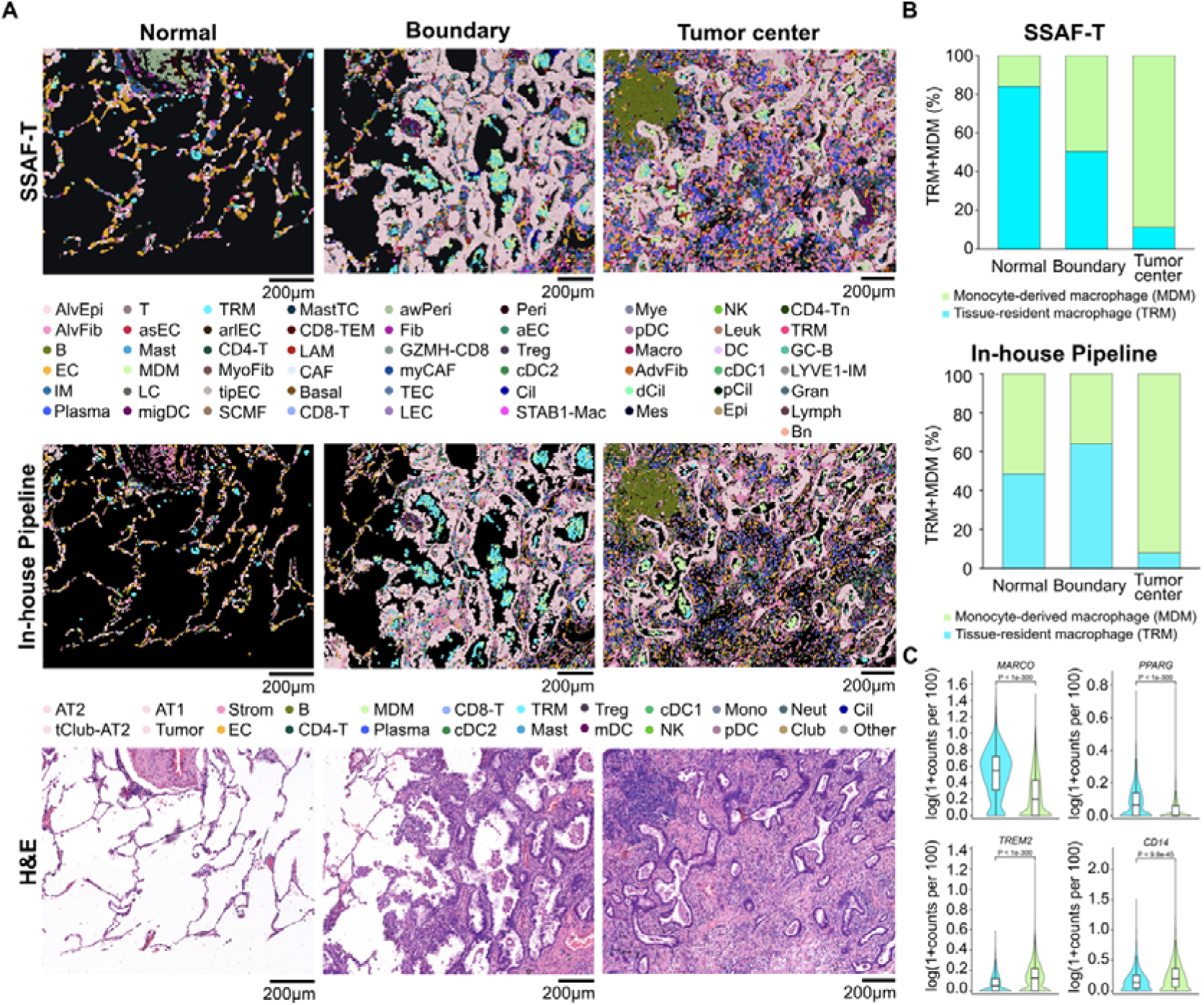
Spatial distribution of tissue-resident and monocyte-derived macrophages in human lung adenocarcinoma. (A) Cell-type maps generated by SSAF-T (top) and an in-house pipeline using Xenium default segmentation followed by SingleR annotation with the LuCA reference atlas (middle), and corresponding H&E-stained images (bottom) of three representative regions: normal lung, tumor boundary, and tumor center. Scale bars, 200 μm. (B) Proportions of tissue-resident macrophages (TRMs; teal) and monocyte-derived macrophages (MDMs; light green) among the two macrophage populations in normal lung, tumor boundary, and tumor center, as identified by SSAF-T (top) and the in-house pipeline (bottom). In normal lung, TRMs within the alveolar spaces predominate. At the tumor border, TRMs and MDMs accumulate together in the alveolar spaces, and MDMs are also scattered throughout the interstitium. In the tumor core, TRMs are largely absent from the glandular lumina, which are instead occupied mostly by MDMs. (C) Expression of representative TRM-associated genes (*MARCO* and *PPARG*, top) and MDM-associated genes (*TREM2* and *CD14*, bottom) in TRMs (n = 3,193) and MDMs (n = 9,454) identified by SSAF-T. Only genes detected in at least 10% of cells of both populations and with a Benjamini–Hochberg-adjusted P < 0.05 are shown.

SSAF-T recapitulated the expected gradient. The proportion of TRMs decreased stepwise from 83.9% in normal lung (n = 62) to 50.3% at the tumor border (n = 791) and 11.2% in the tumor core (n = 251) (Fig. 5B, top). The in-house pipeline also captured the scarcity of TRMs in the tumor core (7.1%, n = 1,147). However, the TRM proportion in normal lung (46.6%, n = 133) was lower than at the tumor border (62.1%, n = 1,336), so the pipeline failed to reproduce the TRM predominance expected in normal lung (Fig. 5B, bottom).

We next asked whether the labels assigned by SSAF-T reflect the transcriptional programs of TRMs and MDMs. Casanova-Acebes *et al.* identified TRM– and MDM-associated genes conserved between mouse and human NSCLC. We compared the expression of seven of these genes included in our panel between TRMs (n = 3,193) and MDMs (n = 9,454) across the entire lung tissue (Fig. 5C). The scavenger receptor gene MARCO and the transcription factor gene PPARG were higher in TRMs. The lipid metabolism gene TREM2, the monocyte identity gene CD14, and CCR2 (P = 4.0 × 10 ²) were higher in MDMs. SIGLEC1 (P = 0.07) and STMN1 (P = 0.54) did not differ significantly. All five differentially expressed genes changed in the reported direction. Thus, SSAF-T distinguish the two macrophage populations in a biologically consistent manner, at the level of both spatial distribution and transcription.

### CeLLMan portal provides interactive visualization and supports human refinements of knowledge graph

To enable researchers to explore the knowledge graph used by CeLLMan and examine the literature supporting its cell-type annotations, we developed an interactive web portal. The knowledge graph covers 41 organs each in human and mouse, comprising 1,450 and 1,605 cell types and 2,966 and 2,494 marker genes, respectively (Supplementary Fig. 3A). These marker genes were curated from 6,070 unique PubMed publications for human and 5,823 for mouse (Supplementary Fig. 3B). The web portal (https://cellman.pnucolab.com) provides an interactive cell-type hierarchy organized by species and organ (Supplementary Fig. 3C). Selecting a cell type displays its marker genes, categorized as defining, required, supporting, or negative, together with the PMIDs of supporting publications. Users can also submit comments and suggestions regarding marker-gene annotations through the Issues tab and review updates and changes to the knowledge graph through the Releases tab.

### SSAF-T portal demonstrates scalable analysis and provides dedicated 3D viewer for intereactive exploration

We present the SSAF-T web portal (https://ssaf.pnucolab.com/T), providing interactive access to 129 spatial transcriptomics datasets processed with SSAF-T. The collection includes 70 Xenium, 26 CosMx, 19 MERSCOPE, 9 Visium HD, and 5 Stereo-seq sections across 19 organs, comprising 112 human and 17 mouse sections under normal, diseased, or treated conditions (Supplementary Fig. 4A). These datasets were collected from HISSTA, SPATCH, DeepSpaceDB, 10x Genomics, and STOmics DB, together with the 14 datasets analyzed in this study, including 9 human heart sections and 1 human placenta section. Users can explore cell-type maps, segmented cells, and gene expression in two or three dimensions, and download the processed SSAF-T results as SpatialData Zarr stores (Fig. 3E, Supplementary Fig. 4B, Supplementary Movie 1). The portal also supports loading and visualizing their own SpatialData datasets directly from local storage without uploading them to a server. We further developed a standalone SSAF-T viewer application with the same visualization capabilities.

### SSAF-T supports large-scale data analyses via pipelined execution

To facilitate automated analysis of spatial transcriptomics data, we integrated SSAF-T with Autopipe^51^, an LLM-based framework for generating and executing reproducible bioinformatics pipelines. This integration allows users to run SSAF-T through natural-language instructions without manually executing individual analysis steps. Users can first configure and refine SSAF-T analysis parameters on a representative dataset, then reuse the resulting pipeline to process multiple samples automatically. This enables consistent application of SSAF-T across multiple spatial transcriptomics samples.

## Methods

### Data acquisition

For spatial transcriptomics benchmarking, we used four publicly available datasets with published cell-type annotations: Visium HD mouse small intestine from Kamel *et al.*, Stereo-seq mouse brain from Chen *et al.*, Xenium mouse kidney from Xuanyuan *et al.*, and MERFISH mouse liver from Liu *et al.* For segmentation-free analysis, we used two additional publicly available Xenium datasets with published cell segmentation and cell-type annotations: human heart from Kuznetsov *et al.* and human placenta from Xenakis *et al.* For case studies, we generated two in-house Xenium datasets: wounded mouse skin collected on post-wounding day 15 and human lung adenocarcinoma tissue profiled using the Xenium Prime 5K human panel. For CeLLMan benchmarking, we used Tabula Sapiens v2.0 and Tabula Muris Senis, retaining the original cell-type annotations. For the SSAF-T web portal, we collected 115 additional spatial transcriptomics datasets from five sources: 87 from HISSTA (46 Xenium, 23 CosMx, and 18 MERFISH), 15 from SPATCH (6 Visium HD, 3 Stereo-seq, 3 Xenium, and 3 CosMx), 10 from DeepSpaceDB (Xenium), 2 from 10x Genomics (Visium HD), and 1 from STOmicsDB (Stereo-seq).

### Generation of the gene expression vector field

For array-based data, gene counts are arranged on a regular spatial grid. Each gene-specific count image is smoothed by separable Gaussian convolution, applied sequentially along the spatial axes. For imaging-based data, spatial gene expression densities are estimated from transcript coordinates using kernel density estimation (KDE) evaluated on a regular grid. The kernel is selectable, with all supported kernels scaled to the same per-axis variance to maintain a consistent spatial bandwidth across kernel types and dimensionalities. For three-dimensional data, KDE is performed over all three spatial axes. At each grid position, the estimated expression densities of individual genes are stacked into a gene expression vector. The L1 norm of each vector is calculated and stored separately as the local molecular density. Each expression vector is then L1-normalized and log1p-transformed. The resulting gene expression vector field is stored in chunked arrays.

### Sampling and clustering of representative vectors

The staining image is first converted to grayscale, and a binary tissue mask is generated by calculating and applying Otsu’s threshold. Spatial positions are sampled uniformly at random within the tissue mask, with the number of sampled positions proportional to the mask area. When multiple staining images are used for three-dimensional analysis, sampling is performed independently for each imaging plane. Positions with local molecular density below a threshold derived from the density contribution of a single transcript are removed. The gene expression vectors corresponding to the sampled positions are extracted from the field and stored in an AnnData object containing the normalized expression matrix, spatial coordinates in micrometers, and gene names. Principal component analysis (PCA) is performed on the expression matrix, followed by k-nearest-neighbor (kNN) graph construction and Leiden community detection.

### A literature-grounded cell-type knowledge graph

We constructed a literature-grounded cell-type knowledge graph organized into species-organ-specific modules. Each cell type is represented as a node containing its standardized name, marker genes, and hierarchical position. Cell types shared across organs are stored in a species-specific common module and linked to organ-specific modules through CONTAINS relationships or REFINES relationships when additional organ-specific markers are present. Cell-type hierarchies are defined by IS_A relationships, while DEVELOPS_INTO relationships represent developmental transitions from progenitors to differentiated cell types. Disease-associated cell identities are linked to disease nodes through OBSERVED_IN relationships. Spatial context nodes represent cell-type distinctions based on tissue location, such as zonation, and are linked to their corresponding lineages through CONTEXT_OF relationships. Additionally, HAS_ANCHOR_MARKER relationships identify residual lineage markers that indicate cellular origin despite phenotypic transitions.

Marker genes are classified as defining–cell-type-specific positive markers, required–positive markers shared among sibling cell types, supporting–nonspecific positive markers, or negative. Markers shared by multiple subtypes are assigned to their common parent, leaving subtype-specific markers at lower levels. Marker-gene categories are determined by their distribution across cell types and updated when assignments change.

We developed a knowledge graph construction agent that reads published studies characterizing cell types and their marker genes to extract cell-type identities, hierarchical relationships, and marker genes. The agent standardizes cell-type names and incorporates the extracted information into the graph, linking each biological relationship to supporting PubMed identifiers (PMIDs). The resulting graph is validated for marker specificity, hierarchical consistency, PMID support, and duplicate identities. The knowledge graph is stored in a Neo4j database and exposed through a read-only HTTP API (https://cellman.pnucolab.com).

### CeLLMan

CeLLMan consists of a conversational LLM that interacts with users, an annotation agent, and a script that computes cluster expression profiles. The annotation agent follows decision rules defined in a skill document. Based on species and organ information from the user, CeLLMan queries the corresponding knowledge graph module through the cellman-kg Model Context Protocol (MCP) server. Annotation proceeds through iterative rounds of hierarchical classification, beginning with broad cell classes and progressively descending to more specific cell types. In each round, three independent annotation agents evaluate the molecular evidence against the knowledge graph and propose cell-type assignments. Their consensus decision determines the cell-type assignment and whether to proceed to the next level of the hierarchy. This process continues until no further refinement is supported by the available marker-gene evidence.

For each cluster, CeLLMan calculates gene-level statistics from the log-normalized expression matrix, including mean expression within the cluster, across other clusters, and across the dataset, the percentage of expressing cells, and the area under the receiver operating characteristic curve (AUC) for distinguishing cells in the cluster from all other cells. For each candidate cell type, CeLLMan evaluates positive and negative markers, identifies markers supported by the observed expression and those absent from the dataset, and calculates overall marker support. CeLLMan also records cluster-level information, including cell count, housekeeping-gene expression, parent cluster, tissue context, potential mixed populations, and gene detection rates in other clusters (Supplementary Fig. S1B).

For clusters requiring further separation, CeLLMan constructs an expression matrix using marker genes from candidate subtypes and their ancestors, excluding genes shared by all candidates. If fewer than two marker genes are available, CeLLMan instead constructs the expression matrix using 50 principal components obtained by PCA. Subclustering is then performed using Leiden on a 15-nearest-neighbor graph constructed from the resulting matrix.

CeLLMan supports human-in-the-loop annotation through natural-language instructions. Automatic annotations are saved in the ‘cellman_annotation’ column of the output H5AD file, while annotations from the human-in-the-loop process are stored in a separate ‘cellman_free_annotation’ column, preserving the original automatic annotations. CeLLMan also generates a decision table containing assigned cell types, supporting markers, and annotation rationales, along with an archive of intermediate profiles, decisions, and knowledge graph records used during annotation.

### Cell-type mapping

Cell-type labels assigned by CeLLMan are propagated from the annotated sampled vectors to the entire gene expression vector field by pixel-level factor inference adapted from FICTURE^13^. The annotated sampled vectors serve as reference observations and their assigned cell types as predefined factors. Each voxel is assigned to a single cell type rather than a mixture of cell types. For voxel *i* and cell type *k*, the the assignment score *ψ_ik_* is defined by

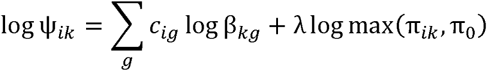

where:

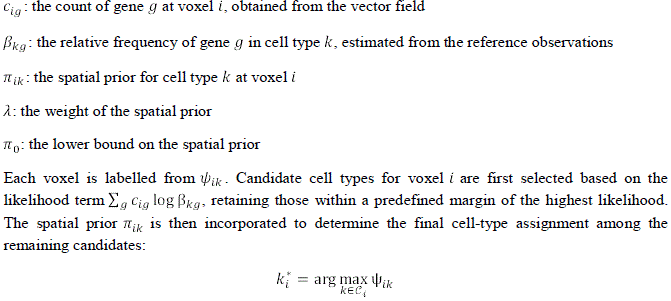

where:

*C_i_*: the set of candidate cell types for voxel *i*

Voxels are subsequently filtered using two criteria: (1) the posterior score 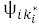 of the assigned cell type and (2) a cell-type-specific molecular density threshold defined as a fraction of that cell type’s maximum density. Voxels falling below either threshold are assigned to the background.

### Segmentation from the cell-type map

Watershed segmentation is performed independently for each cell type, using nuclei seeds and the cell-type map. For two-dimensional data, each seed is assigned the cell-type label at its centroid. For three-dimensional data, the cell-type map is projected along the depth axis to determine the dominant cell type at the each nucleus centroid. The seed is then placed at the median depth of voxels assigned to that cell type along the corresponding vertical column. Segmented objects are filtered using a minimum area threshold. An optional maximum area or volume constraint is applied by retaining pixels or voxels in order of increasing distance from the corresponding seed. Specified cell types can also be excluded from watershed segmentation and processed using connected-component labeling. Each connected region is assigned a single object, and the number of nuclei contained within each object is recorded.

### Benchmarking CeLLMan against published single-cell atlas annotations

We divided Tabula Sapiens v2.0 and Tabula Muris Senis into 29 and 18 organ-specific datasets, respectively, without excluding individual cells. For Tabula Sapiens, the distributed expression matrix, normalized to 10,000 counts per cell and log1p-transformed, was used with the atlas-provided scVI embedding (X_scvi). For Tabula Muris Senis, gene counts were normalized to 10,000 counts per cell and log1p-transformed. The 3,000 highly variable genes were selected separately for each organ using the Seurat method, with assay-specific batch information. The expression matrix was scaled with a maximum value of 10 and reduced to 50 principal components. For organs containing multiple donors or assays, Harmony was applied to correct batch effects. A k-nearest-neighbor graph was constructed using 15 neighbors, followed by Leiden clustering at resolution 1.0 with random seed 0. Preprocessing was performed using Scanpy 1.11.5 and rapids-singlecell 0.13.4 on an NVIDIA RTX 5090 GPU. Each clustered organ dataset was processed using the default annotation mode of CeLLMan, with species and organ information provided as the only contextual inputs. CeLLMan selected the corresponding species-organ module from knowledge graph v1.0. LLM used for annotation was Claude Opus 4.8.

CeLLMan annotations were compared with the published atlas labels using the IS_A hierarchy of the knowledge graph. Each cell was classified into five categories: identical, CeLLMan narrower, CeLLMan broader, conflict, or declined. Cells with non-identical labels were grouped by organ and label pair, and groups containing fewer than 20 cells were excluded from marker-based evaluation. Marker scores were calculated using scanpy.tl.score_genes with marker genes, excluding genes shared by the competing labels. For the CeLLMan narrower and CeLLMan broader categories, ROC AUC was calculated using the marker scores of the narrower cell type to distinguish cells assigned that narrower label from other cells assigned the same parent type. An AUC of 0.65 or higher supported the narrower label, whereas an AUC of 0.55 or lower supported the broader label. For conflicting annotations, marker scores were calculated separately for the CeLLMan label (X) and the published label (Y). The differences between these scores were compared with two reference groups in which both annotations agreed on either X or Y. The marker-score difference was normalized against the two reference groups as follows:

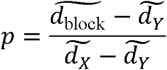

where:

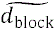: the median of marker-score difference for the evaluated group

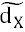: the median of marker-score difference for the reference group in which both annotations agree on cell type X

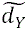: the median of marker-score difference for the reference group in which both annotations agree on cell type Y

A value of p of 0.6 or higher supported CeLLMan, whereas a value of 0.40 or lower supported the published annotation. Intermediate values were classified as undecided.

For each organ, marker-based evaluation outcomes were summarized as percentages of cells within each annotation category. Cells supporting CeLLMan or the published annotation were then aggregated across the CeLLMan narrower, CeLLMan broader, and conflict categories and expressed as percentages of all cells in each organ. These percentages were plotted relative to zero, with positive values indicating support for CeLLMan and negative values indicating support for the published annotation.

### Execution of SSAF-T on spatial transcriptomics datasets

For Visium HD mouse small intestine and Stereo-seq mouse brain, highly variable genes were combined with knowledge-graph marker genes, and Gaussian bandwidths were set to 2.5 and 8 µm, respectively. Five representative vectors were sampled per 10 × 10 µm tissue block, and published nucleus masks were used for segmentation.

For Xenium mouse kidney and MERFISH mouse liver, three-dimensional vector fields were generated with a Gaussian bandwidth of 2.5 µm and 1 µm spatial sampling. Ten representative vectors were sampled per 10 × 10 µm tissue block per imaging plane. Segmentation used vendor-provided Xenium nuclei and Cellpose-detected MERFISH nuclei, respectively.

For Xenium human placenta and human heart, representative vectors were sampled at five and three locations per 10 × 10 µm tissue block, respectively. Nine right-ventricular heart sections (three non-failing, three pressure-loaded, and three failing) were processed independently. Segmentation used vendor-provided Xenium nuclei for placenta and Cellpose-detected nuclei for heart. Syncytiotrophoblast and cardiomyocyte regions were excluded from segmentation and retained as connected structures.

For Xenium mouse skin and Xenium Prime human lung adenocarcinoma, representative vectors were sampled at three and five locations per 10 × 10 µm tissue block, respectively. For mouse skin, tissue masks were generated using 0.6 times the Otsu threshold on the DAPI morphology z-stack. For human lung, imaging planes outside the transcript depth range were excluded. Both datasets used vendor-provided Xenium nuclei for segmentation.

For all datasets, sampled vectors were reduced to 50 principal components, followed by construction of a Euclidean 15-nearest-neighbor graph and Leiden clustering at resolution 1.0.

### Benchmarking SSAF-T on Visium HD mouse small intestine

We reproduced the published ENACT cell segmentation and annotation from Kamel *et al.* following the procedures described in the original study and used them for comparison. Crypt-villus units were identified based on tissue geometry. Units containing at least 20 epithelial cells were selected for analysis. For each unit, the distance of each epithelial cell from the muscularis was measured, and Spearman’s rank correlation coefficient between the distance and the expected epithelial cell-type rank was calculated using ‘scipy.stats.spearmanr’. Epithelial cell-type composition was calculated across all units as the proportion of each label within 10 µm distance intervals from the muscularis, excluding intervals containing fewer than 40 cells.

### Benchmarking SSAF-T on Stereo-seq mouse brain

We used the published cell-type annotations from Chen *et al.* and CellBin segmentation reproduced following the original study’s procedures for comparison, with the bin50 annotation as the reference for cortical layer positions. The pial and white matter boundaries were defined using the ssDNA image and the combined expression of *Mbp*, *Plp1*, and *Mobp*, respectively. The cortex was divided into 250 µm cortical units along the pial boundary, and cortical depth was normalized from the pia to the white matter. For each unit, Kendall’s rank correlation between the expected layer order and median normalized depths was calculated using ‘scipy.stats.kendalltau’. Layer composition was calculated across 20 equal depth intervals. Hippocampal bands were identified from transcript density using Otsu thresholding and assigned to the dentate gyrus, CA1, or CA3 based on marker gene expression. Pearson’s correlation coefficient between the transcript-defined hippocampal bands and the corresponding annotated cell regions was calculated on a 5 µm grid using ‘numpy.corrcoef’. The chance level was estimated from the maximum correlation of 500 random annotation translations of at least 150 µm. CA1 and CA3 were combined for comparison with the published annotation.

### Benchmarking SSAF-T on Xenium mouse kidney

We used the published Baysor cell segmentation and annotation from Xuanyuan *et al.* for comparison. Each published cell was matched to the SSAF-T cell sharing the largest number of transcripts. Segmentation agreement was evaluated using transcript containment, calculated as the number of shared transcripts divided by the total number of transcripts in the published cell, and the Jaccard index, calculated as the number of shared transcripts divided by the total number of unique transcripts across both cells, with both values set to zero for unmatched cells. For runtime comparison, Baysor v0.7.1 was rerun on a 200 µm tissue crop using the authors’ original script and configuration, with their nucleus masks as priors. Execution time for Baysor and SSAF-T was measured as the median wall time of three warm runs on an idle system, excluding agentic annotation from SSAF-T. Transcripts were classified into four groups based on their assignment status in SSAF-T and Baysor. Transcript assignment rates were calculated for each method in 0.5 µm intervals within 12 µm of the nearest nucleus. For each of the four transcript groups, the empirical cumulative distribution function (ECDF) of the distance to the nearest nucleus was calculated. Distances to the nearest nucleus were obtained from the Xenium transcript table. For transcripts excluded by Baysor but recovered by SSAF-T or Xenium default segmentation, multinomial log-likelihood margins were calculated using cell-type reference profiles constructed from the published segmentation. Each margin was defined as the difference between the assigned cell-type log-likelihood per transcript and the highest competing cell-type log-likelihood, with 95% confidence intervals estimated from 200 multinomial bootstrap replicates.

### Benchmarking SSAF-T on Merfish mouse liver

We used the published Cellpose cell segmentation and annotation from Liu *et al.* for comparison. Each published cell was matched to the SSAF-T cell sharing the largest number of transcripts. Segmentation agreement, transcript assignment rates, and empirical cumulative distributions of distance to the nearest nucleus were calculated as described for Xenium mouse kidney. For transcripts excluded by the published segmentation but recovered by SSAF-T, multinomial log-likelihood margins were calculated for seven cell types using reference expression profiles constructed from the published count matrix, with 95% confidence intervals estimated from 300 multinomial bootstrap replicates. The analysis was repeated after excluding knowledge-graph marker genes of the assigned label and of all its ancestors, with the reference profiles renormalized over the remaining genes.

### Segmentation-free analysis

To quantify STB surface contacts, the SSAF-T cell-type map was projected across imaging planes 4-19 using the most frequent nonzero cell-type label at each pixel. STB boundaries were identified using four-neighbor connectivity, and contacts with surrounding cell types were quantified by counting adjacent pixel faces across the tissue. Contact proportions were calculated for each cell type.

For villus composition analysis, villus cores were identified from the three-dimensional cell-type map using six-neighbor connected-component analysis of non-trophoblast villous tissue. STB and CTB voxels within 20 µm were assigned to their nearest villus core. Cell-type volume fractions were calculated and correlated with villus volume using Spearman correlation. Villi were also divided into five equally sized groups by volume, and endothelial fractions were additionally calculated relative to non-STB volume.

Spatial niche analysis was performed on nine Xenium human right-ventricular sections comprising three non-failing (NF), three pressure-overloaded (pRV), and three failing (RVF) hearts. Cardiomyocytes were retained as continuous regions in the SSAF-T cell-type maps, whereas non-cardiomyocyte cells were individually segmented. Analysis positions were defined on a 20 µm grid across tissue regions, and three sets of features were calculated within a 100 µm radius of each position: (i) cardiomyocyte fraction, calculated from the proportion of cardiomyocyte-labeled voxels in the cell-type map; (ii) non-cardiomyocyte composition, calculated from segmented cell centroids grouped into six categories (fibroblasts, endothelial cells, mural cells, myeloid cells, lymphoid cells, and adipocytes); and (iii) non-cardiomyocyte cell density. Cell-type composition was centered-log-ratio transformed with a pseudocount of 1, and cell density was log1p-transformed. Features were standardized across sections and clustered using k-means, fitted on 150,000 randomly sampled positions and applied to all 688,573 positions. The number of niches was selected as the largest k from 3 to 9 satisfying a minimum of 500 positions per niche in at least six sections and a mean leave-one-section-out adjusted Rand index of at least 0.7, yielding eight niches.

### Human lung adenocarcinoma analysis

For comparison with SSAF-T, cells obtained from the default Xenium segmentation were annotated using SingleR^49^ (v2.8.0) with the core lung cancer atlas LuCA^50^ as a reference. Cells with fewer than ten transcripts or fewer than three detected genes were excluded, and transcript counts were normalized to 100 per cell and log-transformed^50^. The reference dataset comprised 266,323 cells from primary tumors profiled using the 10x Genomics platform. These cells were aggregated into pseudo-bulk profiles for each of the 24 major cell types (cell_type_major) using ‘SingleR::aggregateReferencè with default parameters and a random seed of 42, yielding 2,140 reference profiles. Cells with pruned SingleR labels were excluded from subsequent analyses. Following Casanova-Acebes *et al.*, the LuCA labels “Macrophage alveolar” and “Macrophage” and the CeLLMan labels “FABP4-high alveolar macrophage” and “alveolar macrophage” were designated as tissue-resident macrophages (TRMs) and monocyte-derived macrophages (MDMs), respectively. These classifications were based on transcriptional annotations rather than lineage tracing. Because LuCA lacks a distinct interstitial macrophage label, its “Macrophage” population may also include interstitial macrophages. Three regions of interest representing normal lung (1,197 × 996 µm), tumor boundary (1,528 × 1,271 µm), and tumor center (1,785 × 1,367 µm) were selected from the morphology image using Xenium Explorer. Cells were assigned to each region based on their centroids. Within each region, the proportion of TRMs among TRMs and MDMs was calculated separately for SSAF-T and default Xenium segmentation. For the default segmentation, these two populations encompassed all macrophages, whereas SSAF-T cells assigned to other CeLLMan macrophage subtypes were excluded from this comparison.

To evaluate macrophage-associated gene expression, we selected genes reported by Casanova-Acebes *et al.* to be shared between mouse and human TRMs or MDMs in non-small cell lung cancer. Of the 11 reported genes, seven were included in the Xenium panel: MARCO, SIGLEC1, STMN1, and PPARG for TRMs, and TREM2, CCR2, and CD14 for MDMs. Gene expression was compared between TRMs and MDMs across the entire analyzed lung tissue, encompassing both primary tumor pieces rather than only the three selected regions, using the same filtering and normalization procedures. Statistical significance was assessed using a two-sided Wilcoxon rank-sum test (‘scipy.stats.mannwhitneyù), with P values adjusted for multiple testing across the seven genes using the Benjamini–Hochberg method (‘statsmodels.stats.multitest.multipletests’). Individual cells were treated as the unit of analysis. As all cells originated from a single patient, the resulting P values reflect within-sample differences rather than interpatient variation.

### CeLLMan web portal

The CeLLMan web portal is implemented with a SvelteKit (v2.70.3; Svelte v5.56.10) and TypeScript (v6.0.3) frontend and a FastAPI (v0.128.0) backend running on Python 3.12, and each component was containerized and deployed with Docker Compose. The CeLLMan knowledge graph, stored in a Neo4j graph database, is queried only through a separate read-only API, and the backend caches its responses for 30 minutes. For the supporting-literature PMIDs provided by CeLLMan, the web server retrieves bibliographic information, including authors, journal and publication year, from NCBI E-utilities and displays it in the portal. A background worker container runs daily at 12:00 PM KST, compares the current data with the previous snapshot, and records additions, deletions and modifications of marker-gene evidence. The cell-type hierarchy graph was laid out with d3-force (v3.0.0), and all visualizations were rendered as SVG.

### SSAF-T web portal

The SSAF-T web portal was developed using SvelteKit (v2.70.3), Svelte (v5.57.0), and TypeScript (v6.0.3), with a Python 3.11 backend for reading SpatialData (v0.7.3) Zarr stores. The frontend was built using Vite (v8.3.0) and deployed as static files, with the portal and backend served behind an HTTPS reverse proxy using Caddy. The viewer uses WebGL2 for interactive two-dimensional and three-dimensional visualization of cell-type maps, segmented cells, staining images, and transcript coordinates. Large spatial datasets are processed into compressed, spatially indexed tiles and cached for on-demand rendering, with GPU memory management and background workers used to maintain interactive performance.

The published dataset catalog was preprocessed into JSON manifests and cached visualization data. SSAF-T results are provided as downloadable ZIP archives containing SpatialData Zarr stores. The viewer also supports direct loading of local SpatialData Zarr stores through the browser’s File System Access API, without transferring data to the server. Zarr arrays and Parquet-based cell annotations, polygons, and transcripts are read locally, with data processing and surface generation performed using Web Workers.

### SSAF-T viewer application

The SSAF-T viewer is a desktop application for visualizing SSAF-T results stored in Zarr format in 2D and 3D. The application was implemented with Electron, and data processing is handled by a Python server that was packaged into the application with PyInstaller and runs locally when the application starts. When an SSAF-T result is opened, the server reads the structure of the file, determines the layers to display and their spatial placement, and prepares the data for visualization. Data preparation is performed on the CPU and rendering on the GPU with WebGL2. Prepared data are cached and reused when the same dataset is reopened. Installers for Windows and macOS are available from the Downloads tab on the SSAF-T website.

### Mouse wound model

All animal procedures were approved by the Institutional Animal Care and Use Committee of Seoul National University Hospital, Korea (approval no. 22-0166-S1A1) and performed under isoflurane anesthesia. A large full-thickness wound was created in a 7-week-old C57BL/6 mouse by excising a 1.5 cm × 1.5 cm area of dorsal skin. On post-wounding day 15, a skin sample was collected from the wound site, fixed in formalin, and embedded in paraffin. The paraffin-embedded sample was cut into 5-μm sections for analysis using a Xenium Analyzer at Macrogen (Seoul, Korea). After deparaffinization and permeabilization, transcripts were detected with a custom panel targeting 50 skin marker genes, supplemented with the 247-gene 10x mouse brain panel (Supplementary Table 3). Sections were stained with H&E after Xenium analysis. The original Xenium Analyzer output was reprocessed using Xenium Ranger (v2.0), with nuclear boundary expansion set to 2 μm.

## Discussion

In this study, we developed SSAF-T, a framework that identifies cell types directly from spatial gene expression before cell segmentation. In array-based datasets, SSAF-T reconstructed spatial cell-type distributions consistent with known tissue anatomy, while in imaging-based datasets, it produced segmentation results comparable to those of existing segmentation methods, with faster processing times and additional transcript assignments beyond the published segmentations. Based on this, we demonstrated the applicability of SSAF-T across spatial transcriptomics technologies with different spatial resolutions and data characteristics.

An important component of SSAF-T is CeLLMan, an agentic annotation framework that assigns cell-type identities using a literature-grounded knowledge graph. Our benchmarking on single-cell atlases showed that CeLLMan could identify cell types beyond published annotations, demonstrating its potential to complement existing annotation. CeLLMan combines curated marker-gene evidence with expert-guided refinement, allowing researchers to review and refine cell-type annotations. Furthermore, by assigning cell-type identities based on existing biological knowledge, CeLLMan can retain clusters with insufficient evidence as unknown, providing researchers with candidates for further investigation. However, its performance depends on the completeness and accuracy of the knowledge graph. We plan to regularly update and expand the knowledge graph through an automated process with human-on-the-loop validation to incorporate new findings from the literature and improve its coverage of cell types and marker genes.

For all analyses demonstrated in this paper, we used a nuclei-guided watershed method constrained by the SSAF-T cell-type map, considering the absence of a universally accepted gold standard method for cell segmentation. Surprisingly, despite using a relatively basic watershed algorithm, we showed that SSAF-T already produced segmentation results comparable to those of Baysor. Considering that the performance of nuclei-guided watershed segmentation depends on the accuracy of nuclei segmentation methods, adaptation of a better nuclei segmentation method or even a segmentation algorithm without needing nuclei segmentation would potentially further increase the accuracy of cell segmentation in SSAF-T. As improved segmentation methods become available, we plan to extend SSAF-T to support their integration with the segmentation-free cell-type map.

Together, these findings demonstrate that SSAF-T can identify spatial cell-type distributions independently of prior cell segmentation while achieving results comparable to existing segmentation methods on the evaluated metrics. By integrating segmentation-free cell-type mapping, knowledge-grounded automated annotation, and cell segmentation into a single framework, SSAF-T reduces dependence on predefined cell boundaries and reference datasets. This approach provides a flexible basis for spatial transcriptomics analysis, supporting both cell-level and segmentation-free investigations of tissue organization.

## Acknowledgements

During the preparation of this manuscript, the authors used AI-based language tools for the purposes of improving readability and refining the language of the text. The authors also used AI-based coding assistants to aid in the development and debugging of code. The authors reviewed and edited all AI-generated output and took full responsibility for the content of this publication.

## Conflict of Interest

Y.J. and J.P. filed a patent application based on the method discussed in this study. The other authors declare no conflict of interest.

## Availability of data and materials

The source code of SSAF-T and the code to reproduce the results are available at https://github.com/pnucolab/SSAF-T.git. This repository includes CeLLMan as a submodule, linked to its repository(https://github.com/pnucolab/CeLLMan.git).

All datasets analyzed in this study are publicly available. The Tabula Sapiens v2 and Tabula Muris Senis datasets are available from CZ CELLxGENE Discover at https://cellxgene.cziscience.com/collections/e5f58829-1a66-40b5-a624-9046778e74f5 and https://cellxgene.cziscience.com/collections/0b9d8a04-bb9d-44da-aa27-705bb65b54eb, respectively.

The Xenium mouse kidney dataset, including the published Baysor segmentation, is available from GEO under accession GSE269719 (sample GSM8325616). The MERFISH mouse liver dataset, including transcript coordinates, cell boundaries, and cell-type annotations, is available at https://figshare.com/projects/MERFISH_mouse_comparison_study/134213. The Visium HD mouse small intestine dataset is available from 10x Genomics at https://www.10xgenomics.com/datasets/visium-hd-cytassist-gene-expression-libraries-of-mouseintestine. The Stereo-seq adult mouse brain dataset is available from STOmicsDB under accession STDS0000058 (https://db.cngb.org/stomics/mosta/download/).

The Xenium human placenta dataset is available from GEO under accession GSE330886 (sample GSM9734766), with the published annotated Seurat object available at https://doi.org/10.5281/zenodo.20543857. The Xenium human heart dataset is available from GEO under accession GSE345643 (samples GSM10012170, GSM10012171, GSM10012174, and GSM10012175), with the accompanying processed data available at https://doi.org/10.5281/zenodo.20115563.

The raw data for the 115 additional spatial transcriptomics datasets included in the SSAF-T web portal are publicly available from their sources: HISSTA, SPATCH, DeepSpaceDB, 10x Genomics, and STOmicsDB. The SSAF-T-processed datasets, including cell-type maps, cell segmentation results, and gene expression data, are publicly accessible through the SSAF-T web portal (https://ssaf.pnucolab.com/T), where they can be interactively explored and downloaded as SpatialData Zarr files. Xenium human lung adenocarcinoma dataset and Xenium wounded mouse skin dataset are available upon request.

## Authors’ contributions

Y.J. developed SSAF-T, CeLLMan, the CeLLMan knowledge graph, and the SSAF-T web portal, and performed all analyses presented in this study. Ha.L., O.K., and J.-I.K. produced Xenium wounded mouse skin dataset. H.-M.K. developed the SSAF-T viewer application and the AutoPipe pipeline. S.-H.K. and S.P. developed the CeLLMan web portal. He.L, J.B. provided critical comments and contributed to discussions. E.S.J. analyzed Xenium wounded mouse skin dataset. D.L and Y.-L.C analyzed Xenium human adenocarcinoma dataset. J.P. supervised the study.

## Abbreviations

Abbreviations for the cell-type labels used in this study are provided in Supplementary Tables 1 and 2.

## Notes

### Funding

- National Research Foundation of Korea
- National Research Foundation: 2024-00439078)
- Korea Institute for Advancement Technology: RS-2025-02214034
- National Research Foundation: RS-2026-25521031
- Rural Development Administration: RS-2026-25520413

