## Supplementary Figure, Table for "Agentic AI enables cell segmentation-free cell-type inference and cell-type-guided cell segmentation in spatial transcriptomics"

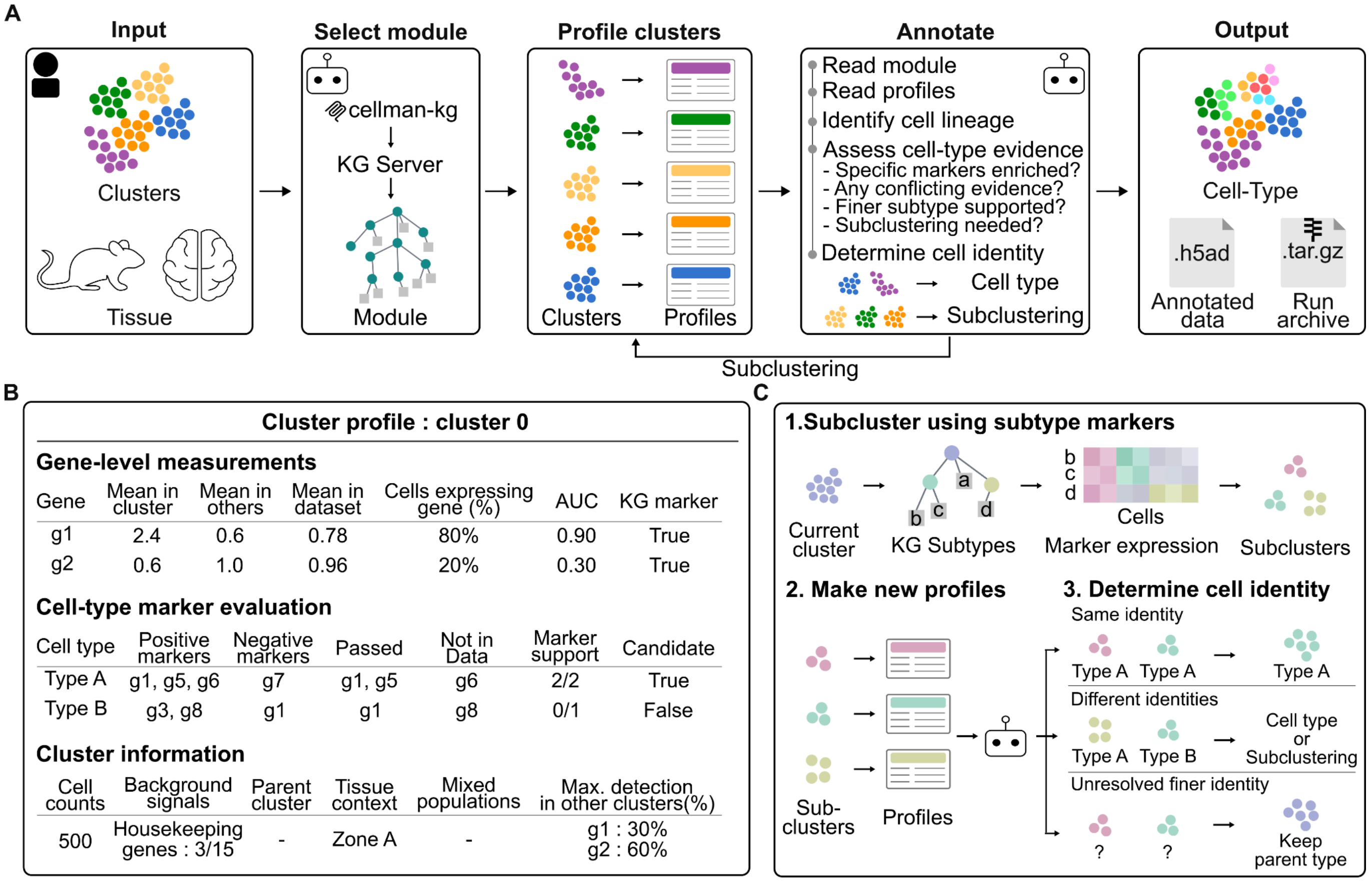


**Figure S1. The workflow of CeLLMan cell-type annotation.** (A) Overview of the "Default Annotation" workflow. CeLLMan takes clustered expression data and tissue information from the user, selects the corresponding module from the CeLLMan knowledge graph through the cellman-kg MCP server, and evaluates each cluster's expression profile against the module's marker genes. Clusters containing more than one cell type are divided into subclusters and re-evaluated. The workflow outputs annotated data in H5AD format and an archive of the annotation run. (B) Cluster expression profiles used for annotation. Each cluster profile includes gene-level statistics, knowledge-graph marker status, and cluster context. (C) Marker-gene-guided subclustering and cell-type annotation. Clusters are re-clustered using marker genes of candidate subtypes from the knowledge graph, and each subcluster is re-evaluated. Subclusters sharing the same identity are merged, those with distinct identities are annotated separately, and unresolved subclusters retain their broader parent cell-type labels.

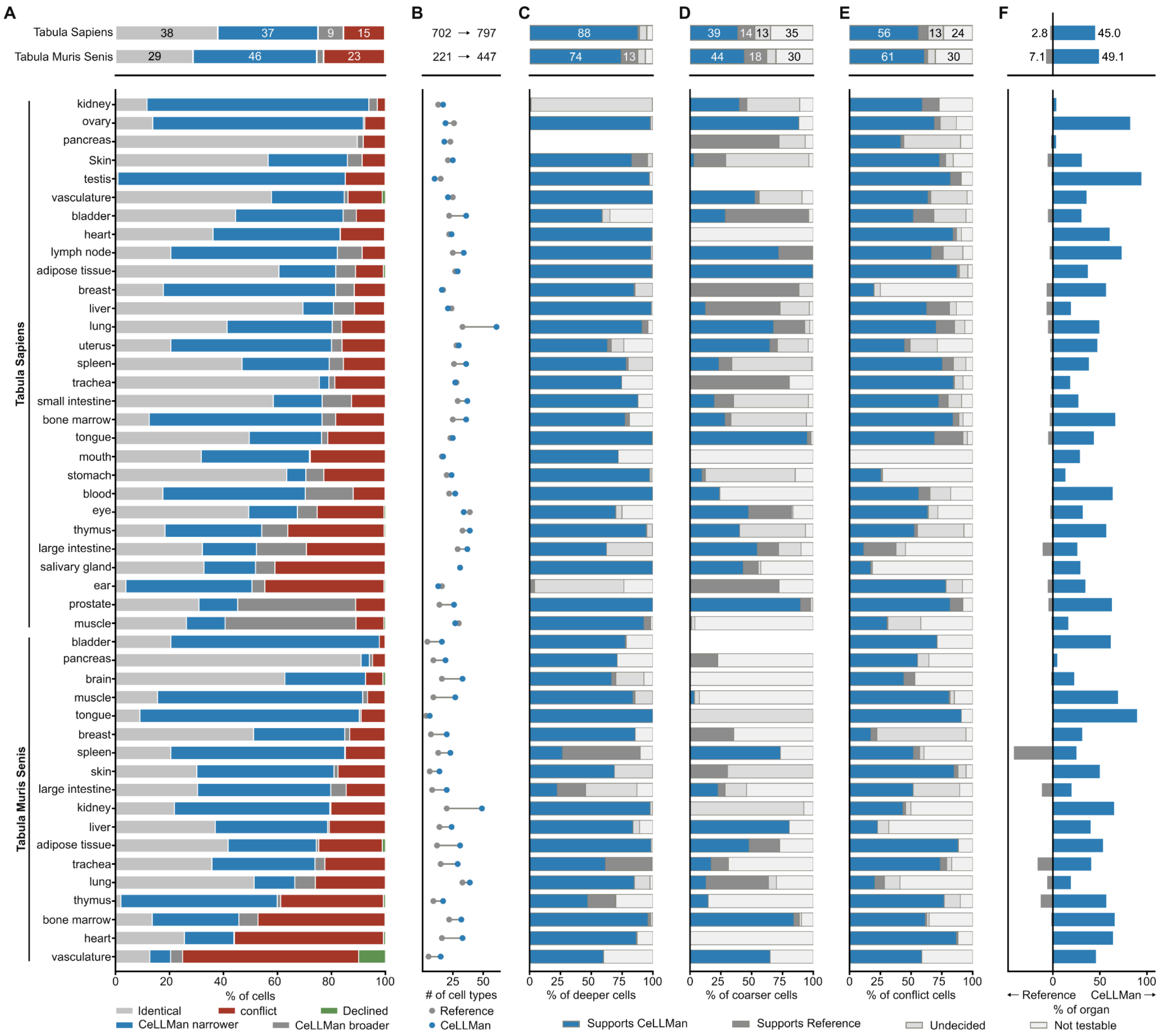


**Figure S2. Benchmarking CeLLMan against the published annotation of Tabula Sapiens and Tabula Muris Senis.** Comparison of CeLLMan's label with the atlas authors' label for the same cell, across 47 organ datasets – Tabula Sapiens (29 organs) and Tabula Muris Senis (18 organs). Above the black line, each atlas as a whole, all of its organs combined into one bar; below the black line, one row per organ, the Tabula Sapiens (A) Stacked bars classifying CeLLMan's label against the author's label into five relations: identical, the same term; CeLLMan narrower, CeLLMan resolved the cell more finely; CeLLMan broader, the authors resolved it more finely; conflict, the two disagree outright, naming unrelated cell types; declined, no label returned. Bar length, percentage of cells. (B) Dumbbell plot of the number of distinct cell types used, the author's annotation (grey) and CeLLMan's (blue). (C-E) Stacked bars of the marker-based assessment of the cells that are not identical. (C) CeLLMan narrower, (D) CeLLMan broader, (E) conflict. Bar length, percentage of cells of that relation; blue, the markers support CeLLMan; dark grey, they support the author; light grey, undecided; white, not testable. (F) Diverging bars of the percentage of each organ's cells for which the markers support CeLLMan's label (right, blue) or the author's (left, grey).

| Abbreviation | Cell type | Abbreviation | Cell type |
|---|---|---|---|
| AA-B | age-associated B cell | LSEC | liver sinusoidal endothelial cell |
| aCap | aerocyte | lSpc | late primary / secondary spermatocyte |
| Acinar | acinar cell | LumSec | luminal secretory cell |
| aCM | atrial cardiomyocyte | Lung-Baso | lung-resident basophil |
| Adipo | adipocyte | Ly6clo-Macro | Ly6c-low infiltrating macrophage |
| AdReg | adipogenesis-regulatory cell | Lym | lymphocyte |
| advFib | adventitial fibroblast | LYVE1-Macro | LYVE1+ tissue-resident macrophage |
| aEC | arterial endothelial cell | LZ-Spc | leptotene-zygotene spermatocyte |
| aEnt | absorptive enterocyte | M | M cell of gut |
| Alpha | pancreatic A cell | mAcinar | mucous acinar cell |
| Alv-Fib | alveolar fibroblast | Macro | macrophage |
| Alv-Macro | alveolar macrophage | MAIT | mucosal-associated invariant T cell |
| Alv-MyoFib | alveolar myofibroblast | Mam-Epi | mammary epithelial cell |
| amNeut | antimicrobial neutrophil | Mam-Lum | mammary luminal epithelial cell |
| apAo-EC | atheroprone aortic endothelial cell | Mam-MPFib | mammary mesenchymal progenitor fibroblast |
| APM | arrector pili smooth muscle cell | MantleB | mantle zone B cell |
| aSat | activated satellite cell | Mast | mast cell |
| ASM | airway smooth muscle cell | MC-TC | tryptase-chymase mast cell |
| ASPC | adipose stem and progenitor cell | mDC | mature dendritic cell |
| Astro | astrocyte | Mel | melanocyte |
| AT0 | alveolar type 0 cell | memB | memory B cell |

| | | | |
|---|---|---|---|
| AT1 | alveolar type 1 cell | memCD8-T | memory CD8 T cell |
| AT2 | alveolar type 2 cell | MEP | megakaryocyte-erythroid progenitor |
| atrGrC | atretic granulosa cell | Mes | mesenchymal cell |
| aw-Fib | airway fibroblast | Mesang | mesangial cell |
| B | B cell | Meso | mesothelial cell |
| Basal | basal cell | MFOL | myelin-forming oligodendrocyte |
| Baso | basophil | mGrC | mural granulosa cell |
| bDuct | basal duct cell | MHCIIhi-PV-Macro | MHCII-high perivascular macrophage |
| Bergmann | Bergmann glia | Micro | microglial cell |
| BEST4-Ent | BEST4+ enterocyte | migDC | migratory dendritic cell |
| Beta | type B pancreatic cell | MK | megakaryocyte |
| BFU-E | BFU-E erythroid progenitor | mNeut | mature neutrophil |
| Bipolar | bipolar cell | MNP | mononuclear phagocyte |
| bKera | basal keratinocyte | Mo-EEC | type Mo enteroendocrine cell |
| BM-mNeut | mature bone marrow neutrophil | moDC | monocyte-derived dendritic cell |
| BMCP | Basophil/Mast cell progenitor | mOligo | mature oligodendrocyte |
| BMEC | brain microvascular endothelial cell | Mono | monocyte |
| Br-Mono | brain monocyte | MPP | multipotent progenitor |
| Brunner | brunner's gland cell | MPP3 | multipotent progenitor 3 |
| bUro | basal urothelial cell | Mrc1-Macro | Mrc1-high resident macrophage |
| capEC | capillary endothelial cell | mregDC | LAMP3+ mature/migratory dendritic cell (mregDC) |

| | | | |
|---|---|---|---|
| CAR | mesenchymal stromal cell / CXCL12-abundant reticular | MS-Macro | medullary sinus macrophage |
| Card-CCR2-moMac | CCR2+ MHC-IIhi monocyte-derived cardiac macrophage | MSC | mesenchymal stem cell |
| Card-CD8-T | CD8-positive cardiac T cell | mTAL | medullary thick ascending limb cell |
| Card-Fib | fibroblast of cardiac tissue | mTEC | medullary thymic epithelial cell |
| Card-MHCIIhi-Macro | resident cardiac macrophage CCR2- MHC-II-hi | Muc | mucus secreting cell |
| Card-ncMono | nonclassical cardiac monocyte | MUC5B-Gob | MUC5B-high goblet cell |
| Card-restFib | resting cardiac fibroblast | Muller | Muller glia |
| Card-Schwann | cardiac Schwann cell / neuronal support cell | Mural | mural cell |
| Card-TIMD4-Macro | resident cardiac macrophage CCR2- TIMD4+ | MuSC | skeletal muscle satellite cell |
| Cb-GABA | cerebellar GABAergic neuron | Musc-Macro | muscularis macrophage |
| Cb-Gran | cerebellar granule cell | Muscle | muscle cell |
| CD-Epi | collecting duct epithelial cell | Myel | myeloid cell |
| CD34-advFib | Cd34-positive adventitial fibroblast | MyelLeuk | myeloid leukocyte |
| CD4-SP | CD4 single-positive thymocyte | MyoEpi | myoepithelial cell |
| CD4-T | CD4-positive, alpha-beta T cell | MyoFib | myofibroblast |
| CD4-TRM | tissue-resident memory CD4 T cell | MyoI | type I myonucleus |
| CD56br-NK | CD56-bright NK cell | MyoI/II | hybrid type I/II myofiber |
| CD56dim-NK | CD56-dim NK cell | MyoIIa | type IIa myonucleus |

| | | | |
|---|---|---|---|
| CD8-CTL | CD8-positive, alpha-beta cytotoxic T cell | MyoIIb | type IIb myonucleus |
| CD8-IEL | CD8 intra-epithelial lymphocyte | MyoIIx | type IIx myonucleus |
| CD8-SP | CD8 single-positive thymocyte | Myom-Fib | myometrial fibroblast |
| CD8-T | CD8-positive, alpha-beta T cell | Myom-PV | myometrial perivascular cell |
| CD8-TRM | tissue-resident memory CD8 T cell | MZ-Macro | marginal zone macrophage |
| cDC | conventional dendritic cell | MZB | marginal zone B cell |
| cDC1 | conventional dendritic cell type 1 | N-EEC | N-cell |
| cDC2 | conventional dendritic cell type 2 | nB | naive B cell |
| cGrC | cumulus granulosa cell | nCD4-T | naive CD4-positive, alpha-beta T cell |
| Chief | chief cell | nCD8-T | naive CD8-positive, alpha-beta T cell |
| Chol | cholangiocyte | ncMono | non-classical monocyte |
| Chondro | chondrocyte | NE | neuroendocrine cell |
| Chor-Fib | choroidal fibroblast | NE-mTEC | neuroendocrine medullary thymic epithelial cell |
| Chor-vEC | choroidal venous endothelial cell | Neo-Glut | neocortical glutamatergic neuron |
| Cil | ciliated cell | Neuron | neuron |
| cLEC | subcapsular sinus ceiling lymphatic endothelial cell | Neut | neutrophil |
| CLP | common lymphoid progenitor | NFOL | newly-formed oligodendrocyte |
| Club | club cell | NK | natural killer cell |
| CM | cardiomyocyte | NKT | natural killer T cell |
| cMono | classical monocyte | nmSchwann | non-myelinating Schwann cell |

| | | | |
|---|---|---|---|
| CMP | common myeloid progenitor | NSC | neural stem cell |
| cMyoEpi | contractile myoepithelial cell | Oligo | oligodendrocyte |
| CNT | connecting tubule cell | OLP-Fib | outer lamina propria fibroblast |
| Col13a1-Fib | Col13a1+ matrix fibroblast | OM-DTL | outer medullary descending thin limb cell |
| colAdvFib | collagen-rich adventitial fibroblast | Oocyte | oocyte |
| Colono | colonocyte | OPC | oligodendrocyte precursor cell |
| Conj-Epi | conjunctival epithelial cell | Oral-bEpi | oral epithelial basal / progenitor stem cell |
| Conj-Gob | conjunctival goblet cell | ORS | outer root sheath suprabasal cell |
| Cor-bEpi | corneal basal epithelial cell | OSE | ovarian surface / coelomic epithelium |
| Cor-Epi | corneal epithelial cell | Ov-Peri | adult ovarian pericyte |
| Cor-sEpi | corneal superficial epithelial cell | Pan | Paneth cell |
| Cor-wEpi | corneal wing epithelial cell | Panc-CD8-T | pancreatic CD8 T cell |
| Crypt-Fib | crypt-niche fibroblast | PancEndo | pancreatic endocrine cell |
| cTAL | cortical thick ascending limb cell | papFib | papillary dermal fibroblast |
| Cxcl14-FAP | Cxcl14+ FAP | Parietal | parietal cell |
| D-EEC | type D enteroendocrine cell | pAstro | protoplasmic astrocyte |
| D1-MSN | D1 medium spiny neuron | PC | collecting duct principal cell |
| D2-MSN | D2 medium spiny neuron | pcHep | pericentral hepatocyte |
| dBasal | dorsal basal cell | pcLSEC | pericentral LSEC |
| DC | dendritic cell | pClub | proximal club cell |
| dClub | distal club cell | pColono | proximal colonocyte |

| | | | |
|---|---|---|---|
| dColono | distal colonocyte | PCV-EC | post-capillary venule endothelium |
| DCS | deep crypt secretory cell | pDC | plasmacytoid dendritic cell |
| DCT | distal convoluted tubule cell | PEC | parietal epithelial cell |
| DCT2 | distal convoluted tubule type 2 cell | pEm-Epi | proliferative endometrial epithelial cell |
| Delta | pancreatic D cell | Peri | pericyte |
| dEnt | distal enterocyte | Perich-Fib | perichondrial fibroblast |
| Det-SMC | detrusor smooth muscle cell | PF-Macro | perifollicular macrophage |
| Deut | deuterosomal cell | Pi16-Fib | Pi16+ universal fibroblast |
| DN | double negative thymocyte | Plac8-moMac | Plac8+ monocyte-derived macrophage |
| DN3 | DN3 thymocyte | Plasma | plasma cell |
| DN4 | DN4 thymocyte | Plt | platelet |
| DP | DP thymocyte | Pod | podocyte |
| Dpp4-FAP | Dpp4+ FAP | POM | periotic mesenchyme |
| DTL | descending thin limb cell | Portal-EC | portal endothelial cell |
| Duct | ductal cell | ppHep | periportal hepatocyte |
| Duct-Macro | ductal macrophage | ppLSEC | periportal LSEC |
| EA-EC | efferent arteriole endothelial cell | PR | photoreceptor cell |
| EC | endothelial cell | preAd-Fib | adipo-regulatory/preadipocyte fibroblast |
| ECC | enterochromaffin cell | preB | pre-B cell |
| eCD4-T | effector CD4-positive, alpha-beta T cell | preColono | colonocyte precursor |
| eCD8-T | effector CD8-positive, alpha-beta T cell | preNeut | pre-neutrophil |
| ECL | enterochromaffin-like cell | proB | Pro-B cell |
| eDuct | excretory duct cell | ProEry | proerythroblast |

| | | | |
|---|---|---|---|
| EEC | enteroendocrine cell | ProMono | promonocyte |
| EGC | enteroglial cell | ProMyelo | promyelocyte |
| Em-Cil | ciliated endometrial epithelial cell | Pros-Basal | basal prostate epithelial cell |
| Em-Epi | endometrial epithelial cell | Pros-Club | club prostate epithelial cell |
| Em-Fib | endometrial stromal fibroblast | Pros-Hillock | hillock prostate epithelial cell |
| Em-PV | endometrial perivascular cell | Pros-intFib | interstitial prostate fibroblast |
| emCD4-T | effector/memory CD4+ T cell | Pros-Lum | prostate luminal epithelial cell |
| emCD8-T | effector memory CD8-positive, alpha-beta T cell | Pros-LumA | luminal acinar prostate epithelial cell |
| Endocard | endocardial cell | Pros-LumD | luminal ductal prostate epithelial cell |
| EoBMP | eosinophil/basophil/mast cell progenitor | Pros-peFib | peri-epithelial prostate fibroblast |
| Epen | ependymal cell | PSC | pancreatic stellate cell |
| Epi | epithelial cell | pSpc | pachytene spermatocyte |
| Epicard | epicardial cell | PT | proximal tubule cell |
| ePOM | epithelium-associated periotic mesenchyme | PTC-EC | peritubular capillary endothelial cell |
| Ery | erythroid cell | PTM | peritubular myoid cell |
| EryB | erythroblast | PTS1 | proximal tubule cell S1 |
| EryP | erythroid progenitor | PTS2 | proximal tubule cell S2 |
| eSpd | elongating/elongated spermatid | PTS3 | proximal tubule cell S3 |
| Fib | fibroblast | Pulm-vEC | pulmonary-venous endothelial cell |
| Fil-Kera | filiform papilla keratinocyte | PV-Fib | perivascular fibroblast |
| FoB | follicular B cell | PV-Macro | perivascular macrophage |

| | | | |
|---|---|---|---|
| FOLR2-Macro | tissue-resident FOLR2-positive macrophage | PVL | differentiated perivascular-like cell |
| Fov | foveolar cell | RBC | erythrocyte |
| FRC | fibroblastic reticular cell | Ret-EC | retinal vascular endothelial cell |
| FT-Epi | fallopian tube epithelial cell | Rod | rod photoreceptor |
| FT-Sec | fallopian tube secretory epithelial cell | RP-Macro | red pulp macrophage |
| GABA | GABAergic neuron | RPE | retinal pigment epithelial cell |
| Gamma | gamma cell | rSpd | round spermatid |
| gCap | general capillary endothelial cell | sAcinar | serous acinar cell |
| GCB | germinal center B cell | Schwann | Schwann cell |
| gdT | gamma-delta T cell | SCS-Macro | subcapsular sinus macrophage |
| Germ | germ cell | sDuct | striated duct cell |
| Ghrl | ghrelin cell | SE-Fib | subepithelial fibroblast |
| gKera | granular keratinocyte | Seb-Duct | sebaceous duct cell |
| Glia | glial cell | Sebo | sebocyte |
| Glom-EC | glomerular endothelial cell | Sec | secretory cell |
| Glut | glutamatergic neuron | SecP | intestinal secretory progenitor cell |
| GMP | granulocyte-monocyte progenitor | sEm-Epi | secretory glandular endometrial epithelial cell |
| Gob | goblet cell | SFRP2-Fib | SFRP2/DPP4+ dermal fibroblast |
| Gob2 | goblet-2 cell | SG | salivary gland cell |
| Gon-Fib | gonadal interstitial mesenchyme / stromal fibroblast | Sinus-EC | sinusoidal endothelial cell |
| Granulo | granulocyte | sKera | spinous keratinocyte |

| | | | |
|---|---|---|---|
| GrC | granulosa cell | SkM | skeletal muscle cell |
| GZMB-CD8-T | GZMB+ CD8+ T cell | SMC | smooth muscle cell |
| GZMK-CD8-T | GZMK+ CD8+ T cell | SMG-Muc | submucosal gland mucous cell |
| GZMKneg-NK | CD56dim GZMK- NK cell | SMG-Ser | submucosal gland serous cell |
| HemSC | hematopoietic stem cell | Spc | spermatocyte |
| Hep | hepatocyte | Spd | spermatid |
| HEV | high endothelial venule cell | SPG | spermatogonium |
| HF-Kera | hair follicle keratinocyte | Spl-Ery | splenic erythroid-lineage cell |
| HFSC | bulge hair follicle stem cell | SPP1-Macro | SPP1+ macrophage |
| Hillock | hillock cell | SqEpi | stratified squamous epithelial cell |
| HM | hair matrix | SSC | undifferentiated spermatogonial stem cell |
| HPC | hematopoietic progenitor cell | Str-GABA | striatal GABAergic neuron |
| HS | luminal hormone-sensing cell | Strom-Macro | stromal tissue-resident macrophage |
| HS-AV | HS-AV hybrid luminal cell | Sweat | eccrine sweat-gland secretory cell |
| HSC | hepatic stellate cell | Sys-vEC | systemic-venous endothelial cell |
| Hsd11b1-Fib | Hsd11b1+ cardiac fibroblast | T | T cell |
| HSPC | hematopoietic stem and progenitor cell | TA | transit amplifying cell |
| iB | immature B cell | TAL | thick ascending limb cell |
| IC | intercalated cell | Taste | taste receptor cell |
| ICA | type A intercalated cell | TEC | thymic epithelial cell |

| | | | |
|---|---|---|---|
| ICB | type B intercalated cell | TEMRA | terminally differentiated effector CD8-positive, alpha-beta T cell |
| ICC | interstitial cell of Cajal | Teno | tenocyte |
| IE-Macro | inner ear macrophage | Test-Macro | tissue-resident testicular macrophage |
| IE-Mel | inner ear melanocyte | Th1 | T helper 1 cell |
| IE-Schwann | inner ear Schwann cell | Theca | theca cell |
| IE-Sec | inner ear endolymph-secreting epithelial cell | Thy-capEC | thymic capillary endothelial cell |
| IEC | intestinal epithelial cell | Thy-capsFib | thymic capsular fibroblast |
| IgA-PC | IgA plasma cell | Thy-CD56br-NK | thymic CD56bright NK cell |
| IgG-PC | IgG plasma cell | Thy-CD56dim-NK | thymic CD56dim NK cell |
| IgM-PC | IgM plasma cell | Thy-cDC1 | thymic conventional DC1 |
| iGranulo | immature granulocyte / neutrophil precursor | Thy-ILC1 | thymic group-1 ILC / NK cell |
| ILC | innate lymphoid cell | Thy-intFib | interstitial thymic fibroblast |
| IM-DTL | inner medullary descending thin limb cell | Thy-KIR-NK | thymic KIR+NKG2A+ NK cell |
| imB | immature/mature B cell | Thy-med-Macro | thymic medullary macrophage |
| iMicro | injury-associated microglia | Thy-medFib | medullary interlobular thymic fibroblast |
| iNeut | immature neutrophil | Thy-Mural | thymic mural cell |
| Inmt-Fib | inmt+ cardiac fibroblast | Thy-Myoid | myoid thymic epithelial cell |
| Int-Macro | interstitial macrophage | Thy-pDC | thymic plasmacytoid DC |
| intMono | intermediate monocyte | Thy-SC-Macro | thymic subcapsular macrophage |
| Iono | ionocyte | Thy-Treg | thymic regulatory T cell |
| ISC | intestinal stem cell | Thy-vEC | thymic venous endothelial cell |

| | | | |
|---|---|---|---|
| ISP | immature single-positive thymocyte | Thymo | thymocyte |
| iTheca | internal theca cell | Tong-SkM | tongue skeletal muscle cell |
| iUro | intermediate urothelial cell | TR-Macro | tissue-resident macrophage |
| JG | juxtaglomerular cell | TrabM | trabecular meshwork cell |
| K-EEC | type K enteroendocrine cell | TRB-Sec | terminal and respiratory bronchiole secretory cell |
| Kera | keratinocyte | Treg | regulatory T cell |
| Kerato | keratocyte | TRem-Macro | tissue-remodeling macrophage |
| Kid-Epi | kidney epithelial cell | Troph | trophocyte |
| Kid-MHCIIhi-Macro | resident kidney macrophage (MHCII-high) | Tuft | tuft cell |
| Krt13-Basal | squamous/hillock-like basal cell (Krt13+) | UBC | cerebellar unipolar brush cell |
| KSC | keratinocyte stem cell | uHFSC | upper bulge hair follicle stem cell |
| Kup | Kupffer cell | Umb | umbrella cell |
| L-EEC | type L enteroendocrine cell | uNK | uterine natural killer cell |
| LA-Macro | lipid-associated macrophage | uNK1 | uterine natural killer cell type 1 |
| Lac-Acinar | lacrimal gland acinar cell | uNK3 | uterine natural killer cell type 3 |
| Lac-MyoEpi | lacrimal gland myoepithelial cell | Ureth-Lum | urethral luminal epithelial cell |
| lateProB | late pro-B cell | Uro | urothelial cell |
| LC | Langerhans cell | Ut-EC | uterine blood endothelial cell |
| LEC | lymphatic endothelial cell | Ut-PV | uterine perivascular / pericyte cell |
| LEPC | limbal epithelial progenitor cell | vBasal | ventral basal cell |

| | | | |
|---|---|---|---|
| Leuk | leukocyte | vCM | ventricular cardiomyocyte |
| Leydig | Leydig cell | VDC | vestibular dark cell |
| Ling-bKera | lingual basal keratinocyte | vEC | venous endothelial cell |
| Ling-dKera | lingual differentiated keratinocyte | Ven-EC | venular endothelial cell |
| Ling-LC | lingual Langerhans cell | VIC | valvular interstitial cell |
| Ling-sbKera | suprabasal lingual keratinocyte | VSC | vestibular supporting cell |
| Littoral | splenic littoral endothelial cell | VSMC | vascular smooth muscle cell |
| LL-PC | long-lived plasma cell | wAdipo | white adipocyte |
| LMPP | lymphoid-primed multipotent progenitor | Wif1-Fib | Wif1+ cardiac fibroblast |
| lrNK | liver-resident NK cell | | |

**Table S1. Cell-type abbreviations for the scRNA-seq datasets.** Abbreviations of the 473 cell types in the Tabula Sapiens and Tabula Muris Senis datasets, covering both the CeLLMan and the published annotations.

| Abbreviation | Cell type | Abbreviation | Cell type |
|---|---|---|---|
| ABC | arachnoid barrier cell | L4 IT | layer 4 intratelencephalic glutamatergic neuron |
| aEC | arterial endothelial cell | L5 ET | layer 5 extratelencephalic glutamatergic neuron |
| aEnt | absorptive enterocyte | L6 CT | layer 6 corticothalamic glutamatergic neuron |
| Amy-Glut | amygdalar glutamatergic neuron | L6 IT | layer 6 intratelencephalic glutamatergic neuron |
| Astr1 | astrocyte subtype 1 | L6b | layer 6b glutamatergic neuron |
| Astr2 | astrocyte subtype 2 | Lamp5 | Lamp5+ GABAergic interneuron |
| Astr3 | astrocyte subtype 3 | Leuk | leukocyte |
| Astr4 | astrocyte subtype 4 | LSEC | liver sinusoidal endothelial cell |
| Astr5 | astrocyte subtype 5 | M | M cell of gut |
| Astro | astrocyte | Macro | macrophage |
| B | B cell | Mes | mesenchymal cell |
| BAM | border associated macrophage | Micro | microglial cell |
| BLA | basolateral amygdala principal glutamatergic neuron | Mono | monocyte |
| BMEC | brain microvascular endothelial cell | Mural | mural cell |
| botEnt | villus-bottom enterocyte | Muscle | muscle cell |
| BST | extended amygdala / bed nucleus of the stria terminalis GABAergic neuron | MyoFib | myofibroblast |
| CA1 | hippocampal CA1 pyramidal neuron | Neo-Glut | neocortical glutamatergic neuron |
| CA3 | hippocampal CA3 pyramidal neuron | Neuron | neuron |

| | | | |
|---|---|---|---|
| Cck | CCK+ GABAergic cortical interneuron | Neut | neutrophil |
| CD-Epi | collecting duct epithelial cell | NK | natural killer cell |
| CD4-T | CD4-positive T cell | NKT | natural killer T cell |
| CD8-T | CD8-positive T cell | Oligo | oligodendrocyte |
| Chol | cholangiocyte | OPC | oligodendrocyte precursor cell |
| CNT | connecting tubule cell | Pan | Paneth cell |
| Crypt | crypt cell | PC | collecting duct principal cell |
| cTAL | cortical thick ascending limb cell | pcHep | pericentral hepatocyte |
| Ctx-IN | cortical GABAergic interneuron | pcLSEC | pericentral LSEC |
| DA | dopaminergic neuron | PEC | parietal epithelial cell |
| DCT | distal convoluted tubule cell | Peri | pericyte |
| DCT2 | distal convoluted tubule type 2 cell | pFib | portal fibroblast |
| dEnt | distal enterocyte | Plasma | plasma cell |
| DG | dentate gyrus granule cell | Pod | podocyte |
| DTL | descending thin limb cell | ppHep | periportal hepatocyte |
| EC | endothelial cell | ppLSEC | periportal LSEC |
| ECC | enterochromaffin cell | PT | proximal tubule cell |
| EEC | enteroendocrine cell | PTS1 | proximal tubule cell S1 |
| Ent | enterocyte | PTS2 | proximal tubule cell S2 |
| Epen | ependymal cell | PTS3 | proximal tubule cell S3 |
| Epi | epithelial cell | Pvalb | PVALB+ GABAergic interneuron |
| EX L5/6 | layer 5/6 excitatory neuron | Pvalb-Gad1 | PVALB+ GAD1+ GABAergic interneuron |

| | | | |
|---|---|---|---|
| EX Mb | midbrain excitatory neuron | RBC | erythrocyte |
| Fib | fibroblast | Reg4-ECC | Reg4+ enterochromaffin cell |
| FR_PT | failed-repair proximal tubule cell | S | S-cell |
| GABA | GABAergic neuron | Sec | secretory cell |
| Glia | glial cell | SMC | smooth muscle cell |
| Glom-EC | glomerular endothelial cell | SNc-DA | substantia nigra pars compacta dopaminergic neuron |
| Glut | glutamatergic neuron | Sst | sst+ GABAergic cortical interneuron |
| Gob | goblet cell | Sst-Chodl | sst Chodl long-range GABAergic cortical interneuron |
| Gob-Pan | goblet-Paneth intermediate cell | Str-GABA | striatal GABAergic neuron |
| Hep | hepatocyte | T | T cell |
| HPC-Pyr | hippocampal pyramidal neuron | TA | transit amplifying cell |
| HSC | hepatic stellate cell | TAL | thick ascending limb cell |
| IC | intercalated cell | Telo | telocyte |
| ICA | type A intercalated cell | Th-GABA | thalamic GABAergic neuron |
| ICB | type B intercalated cell | Th-Glut | thalamic glutamatergic neuron |
| ICC | interstitial cell of Cajal | Th-Mtx | matrix thalamic relay neuron |
| IEC | intestinal epithelial cell | tipEnt | villus-tip enterocyte |
| iEnt | immature enterocyte | Troph | trophocyte |
| IgA-PC | IgA plasma cell / plasmablast | Tuft | tuft cell |
| IM-DTL | inner medullary descending thin limb cell | Tuft2 | tuft cell type 2 |

| | | | |
|---|---|---|---|
| Imm | immune cell | Uro | urothelial cell |
| Inj_PT | injured proximal tubule cell | vEC | venous endothelial cell |
| iPan | immature Paneth cell progenitor | Vip | vip+ GABAergic cortical interneuron |
| ISC | intestinal stem cell | visSMC | visceral smooth muscle cell |
| Kup | Kupffer cell | VLMC | vascular leptomeningeal cell |
| L-EEC | type L enteroendocrine cell | VSMC | vascular smooth muscle cell |
| L2/3 IT | layer 2/3 intratelencephalic glutamatergic neuron | CTB | cytotrophoblast |
| EVT | extravillous trophoblast | HBC | Hofbauer cell |
| iEVT | interstitial extravillous trophoblast | Mast | mast cell |
| PVF | placental villous fibroblast / mesenchymal stromal cell | STB | syncytiotrophoblast |
| Adipo | adipocyte | cFib | fibroblast of cardiac tissue |
| CM | cardiomyocyte | EpiC | epicardial cell |
| cDC2 | conventional dendritic cell type 2 | Macro-P | macrophage, proliferating |
| cEC | capillary endothelial cell | Mono-P | monocyte, proliferating |
| cNK | cardiac NK cell | Mye | myeloid cell |
| cSchw | cardiac Schwann cell / neuronal support cell | Mye-P | myeloid cell, proliferating |
| DC | dendritic cell | rcFib | resting cardiac fibroblast |
| LEC | lymphatic endothelial cell | TRM | tissue-resident memory CD8 T cell |
| Lymph | lymphocyte | APM | arrector pili smooth muscle cell |
| KC | keratinocyte | ASPC | adipose stem and progenitor cell |

| | | | |
|---|---|---|---|
| MDM | monocyte-derived macrophage | BK | basal keratinocyte |
| ORS | outer root sheath | Bulge HFSC | bulge hair follicle stem cell |
| PapFib | papillary dermal fibroblast | CMyo | committed myocyte |
| RetFib | reticular dermal fibroblast | DP | dermal papilla |
| SK | spinous keratinocyte | DS | dermal sheath |
| AdvFib | adventitial fibroblast | GZMH-CD8 | gzmh-positive cd8-positive, alpha-beta t cell |
| AlvEpi | alveolar fibroblast | LAM | TREM2-high lipid-associated macrophage |
| arlEC | arteriolar endothelial cell | LC | Langerhans cell |
| asEC | endothelial cell, angiogenic stalk | LYVE1-IM | LYVE1-positive interstitial macrophage |
| awPeri | airway pericyte | MastTC | tryptase-chymase mast cell |
| Basal | basal cell | migDC | mature migratory dendritic cell |
| Bn | naive B cell | myCAF | myofibroblastic cancer-associated fibroblast |
| CAF | cancer-associated fibroblast | pCil | proximal ciliated cell |
| CD4-Tn | naive CD4-positive, alpha-beta T cell | pDC | plasmacytoid dendritic cell |
| CD8-TEM | effector memory CD8-positive, alpha-beta T cell | SCMF | secondary crest myofibroblast |
| cDC1 | conventional dendritic cell type 1 | STAB1-Mac | STAB1+ macrophage |
| Cil | ciliated cell | TEC | tumor endothelial cell |

| | | | |
|---|---|---|---|
| dCil | distal ciliated cell | tipEC | tip endothelial cell |
| GC-B | germinal center B cell | Treg | regulatory T cell |
| Gran | granulocyte | TRM | tissue-resident macrophage |
| AT1 | alveolar type 1 cell | AT2 | alveolar type 2 cell |
| Strom | stromal cell | Club | club cell |
| tClub-AT2 | transitional club/AT2 cell | mDC | mature dendritic cell |

**Table S2. Cell-type abbreviations for the spatial transcriptomics datasets.** Abbreviations of the cell types for the Xenium mouse kidney, MERFISH mouse liver, Stereo-seq mouse brain and Visium HD mouse small intestine datasets, for the Xenium human placenta and Xenium human heart datasets in Figures S5 and S6, and for the Xenium wounded mouse skin and Xenium human adenocarcinoma datasets.

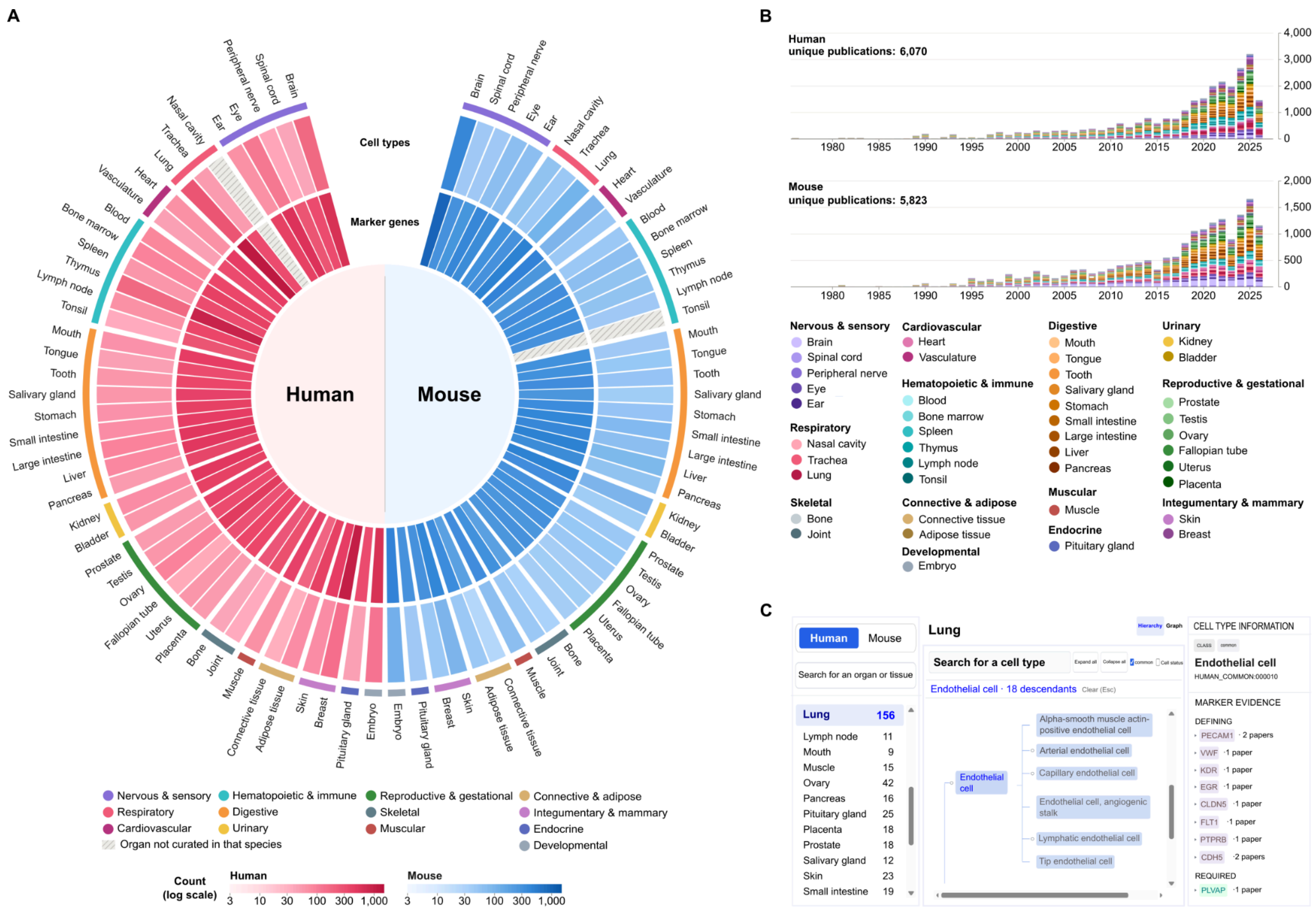


**Figure S3. Overview of CeLLMan web interface.** (A) Number of curated cell types (outer ring) and marker genes (inner ring) for each organ in human (red) and mouse (blue), shown on a log scale. (B) Number of unique PubMed publications per year used for curation in human and mouse, stacked by organ. (C) The CeLLMan web interface. It shows the cell type hierarchy, the marker genes CeLLMan uses for the selected cell type, and the PMIDs supporting each marker.

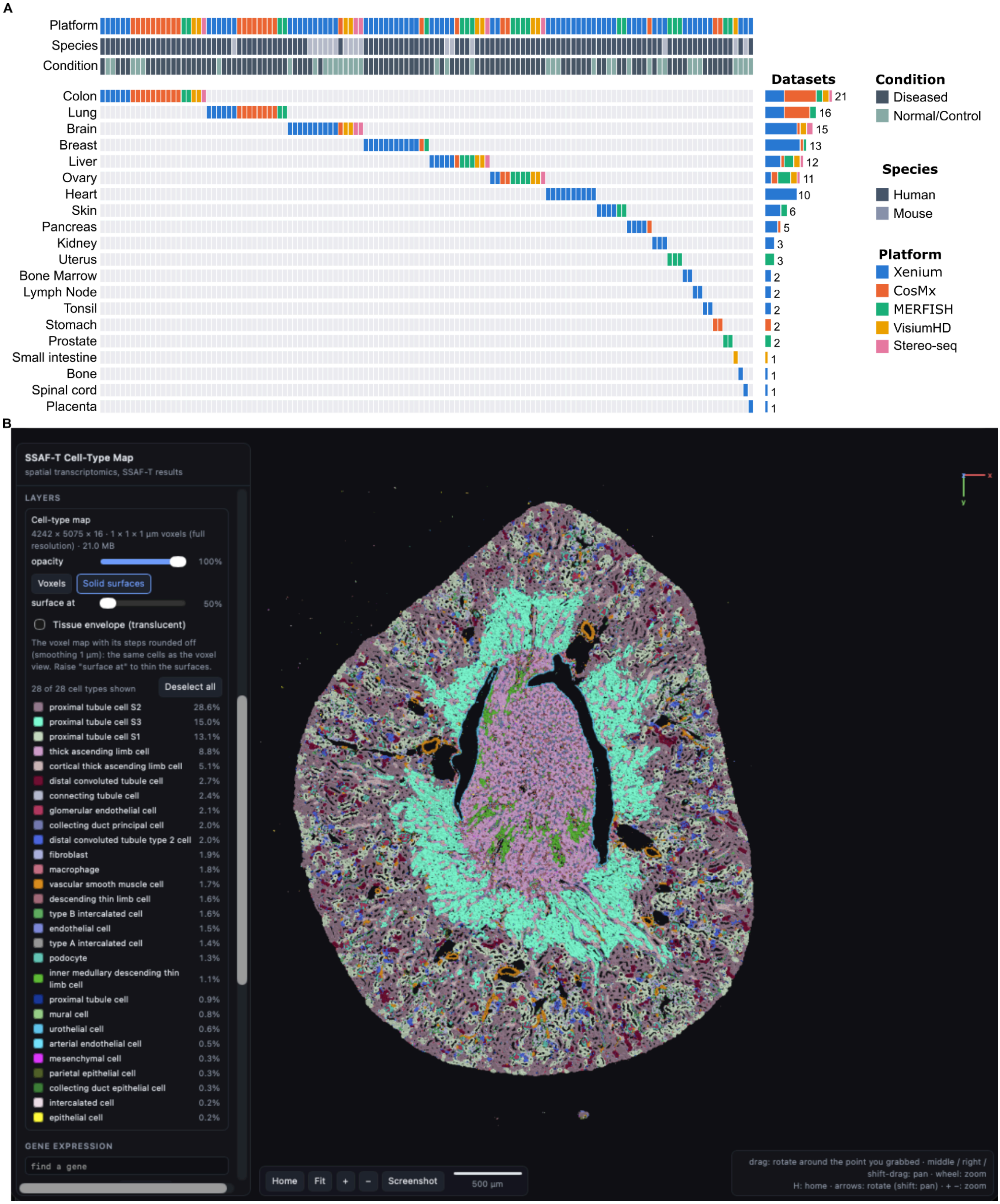


**Figure S4. Overview of the SSAF-T web portal.** (A) Distribution of 129 spatial transcriptomics datasets processed with SSAF-T across organs, platforms, species, and conditions. Each column represents a dataset, with platform, species, and condition indicated by the color bars at the top. Bar plots on the right indicate the number of datasets for each organ. (B) Interactive three-dimensional visualization of the SSAF-T cell-type map in Xenium mouse kidney, showing cell-type-specific spatial distributions and controls for selecting cell types, adjusting visualization settings, and exploring gene expression.

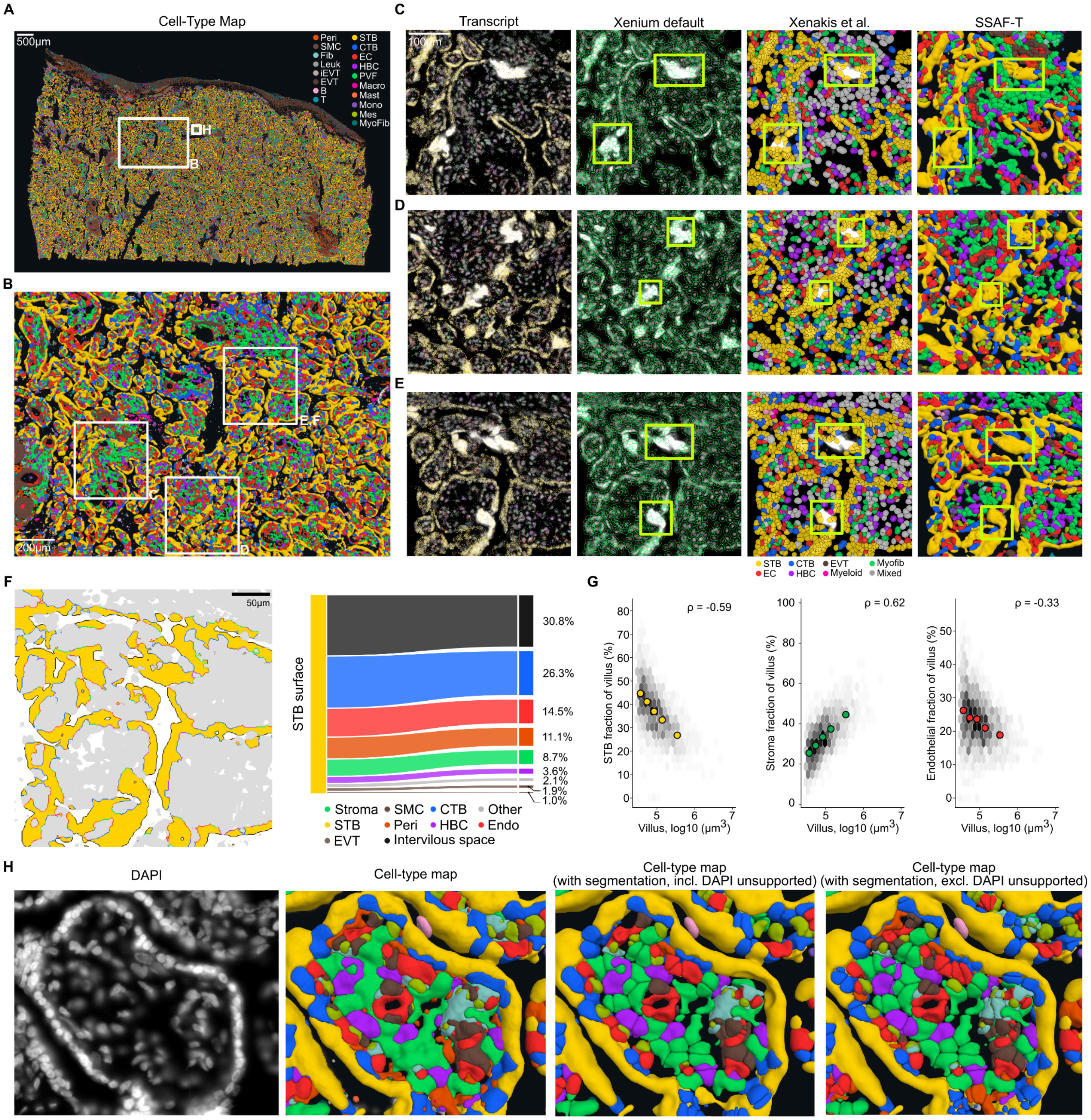


**Figure S5. Segmentation-free analysis for syncytiotrophoblast (STB) on Xenium human placenta.** (A) SSAF-T cell-type map of the whole tissue section. The box indicates the region enlarged in B. (B) Enlarged cell-type map with boxes indicating the regions shown in C-F. (C-E) Comparison of transcript distributions, Xenium default segmentation, published cell-type annotations from *Xenakis et al.*, and SSAF-T cell-type maps in three representative regions. Marker transcripts are shown for STB (*CRH, PAPPA, PLAC4, EBI3*), CTB (*ELF5, MET*), Hofbauer cells (*VSIG4, CD163*), endothelial cells (*PECAM1, VWF, ERG*), and PVF (*THY1, PDGFRA, COL5A2*). Green boxes indicate regions with STB marker expression that are excluded from Xenium default segmentation but identified as STB by SSAF-T. (F) Representative STB territory (yellow) and surrounding cell types (grey), with STB boundaries colored according to the adjacent cell type. Composition of cell types adjacent to STB boundaries across the tissue section, calculated from the SSAF-T cell-type map. (G) Relationship between villus volume and the proportions of STB, stroma, and endothelial cells across approximately 2,300 villi. Grey points indicate villus density, colored points indicate medians of five volume groups, and ρ indicates Spearman correlation. (H) Comparison of DAPI staining, the SSAF-T cell-type map, and cell-type maps after segmentation, either including or excluding DAPI-unsupported regions.

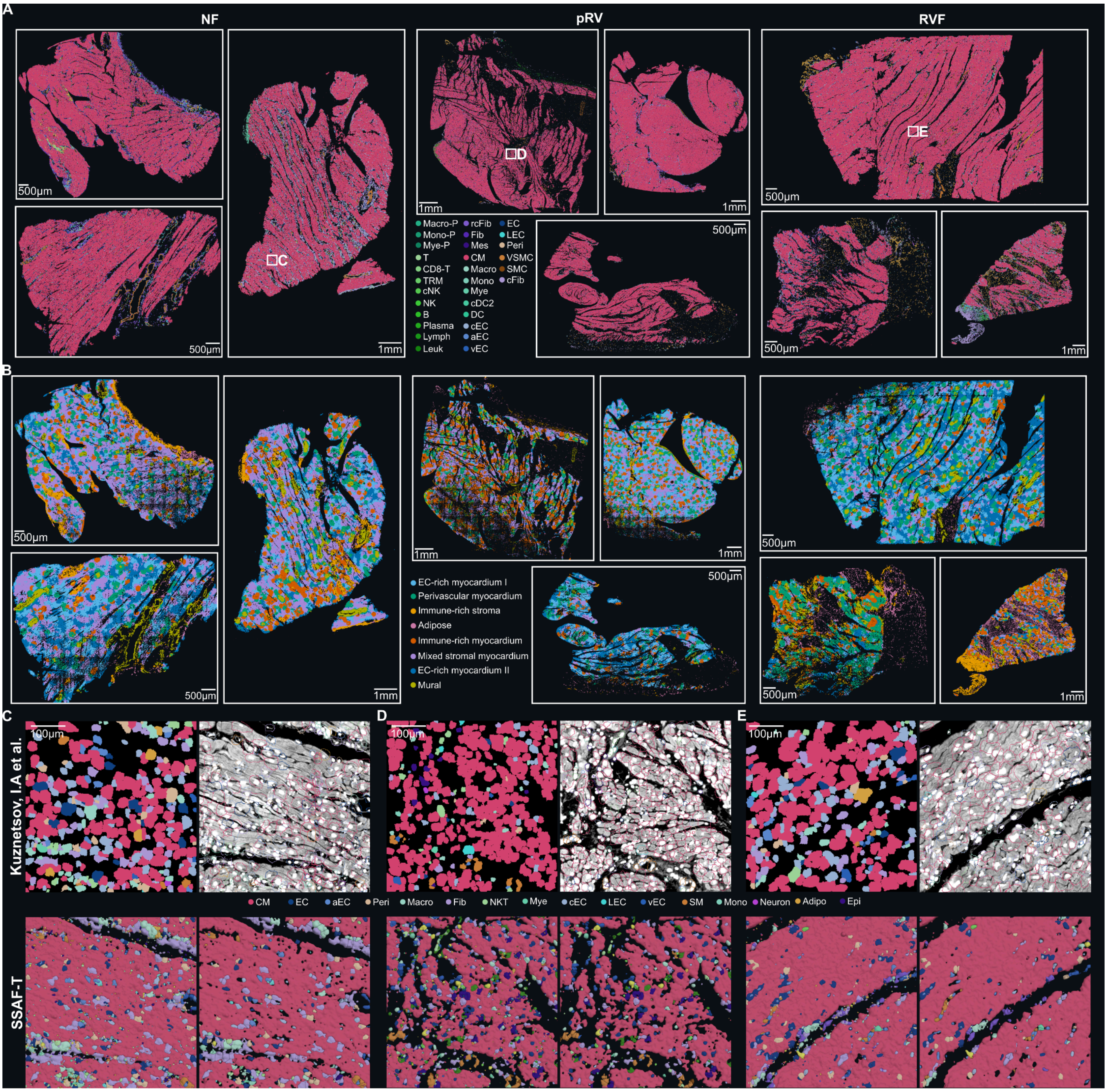


**Figure S6. Segmentation-free analysis for Cardiomyocytes on Xenium human heart.** (A) SSAF-T cell-type maps of nine human right-ventricular sections from non-failing (NF), pressure-overloaded (pRV), and failing (RVF) hearts, with three sections per condition. Cardiomyocytes (pink) were retained as continuous regions without segmentation, whereas non-cardiomyocyte populations were segmented into individual cells. White boxes indicate representative regions enlarged in C–E. (B) Spatial niche maps corresponding to the nine sections in A. Eight niches were identified based on cardiomyocyte area fraction, segmented non-cardiomyocyte cell-type composition, and cell density within a 100 μm radius. Colors indicate distinct niches characterized by their cellular composition. (C–E) Enlarged regions from NF (C), pRV (D), and RVF (E) sections, showing published cell-type annotations and Proseg segmentation from Kuznetsov *et al.* (top) and SSAF-T cell-type maps with partial segmentation (bottom). In the published results, cell-type annotations (left) and Proseg cell boundaries overlaid on DAPI images (right) are shown. SSAF-T results show the corresponding regions with cardiomyocytes retained as continuous structures and non-cardiomyocyte cells individually segmented.

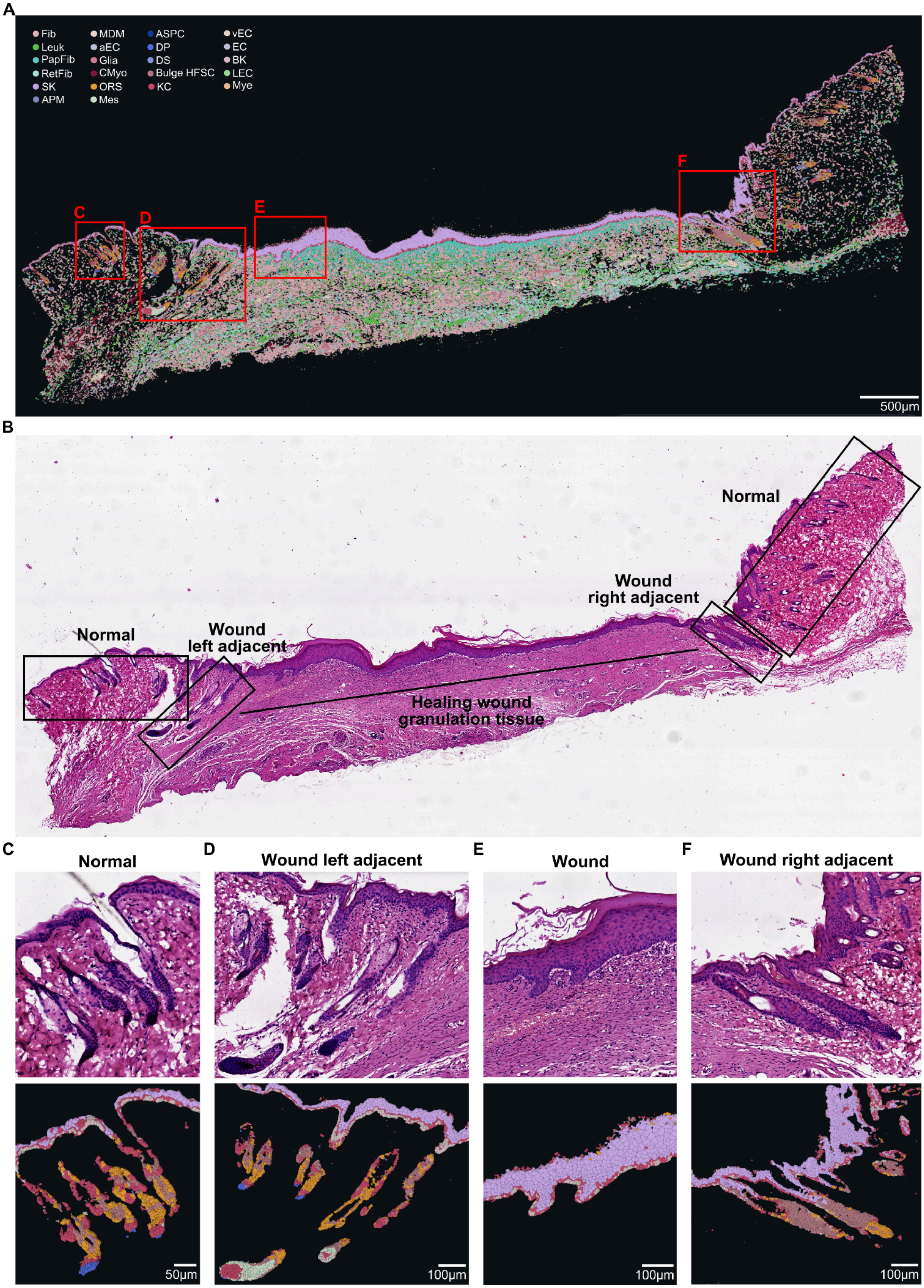


**Figure S7. Appendage like structures observed in the mouse skin wound model.** (A) Overall structure and cell type annotation of the mouse skin sample section. Scale bars are inserted in their relevant image panels. (B) H&E-stained image of the entire tissue section with the corresponding states indicated. (C) Normal telogen phase hair follicles minimally affected by the injury. Multiple telogen hair follicles with attached sebaceous glands, old hair shafts, bulge structures and dermal papillas. (D) Two wound adjacent mid-anagen hair follicles with attached sebaceous glands, the outer root sheath, the matrix core and an unresolved 'mesenchymal cell' labelled cell type layer (colored light green). (E) The healing wound region overlaid with hyperplastic epithelium (white) on granulation tissue. Two epithelial pegs are seen growing downward into the stroma, but lack definite hair follicle components. (F) Reactive epithelial hyperplasia, mimicking hair follicles structure on the right adjacent to the wound. Relatively homogenous epithelial elongations are observed with keratin plugs in the center. There is no evidence of attached appendage glands, dermal papilla, matrix, or sheath organization in these structures.
